# Temporal order separates generic persistence from cue-directed commitment in cell migration

**DOI:** 10.64898/2026.07.16.739021

**Authors:** Subhajit Dutta

## Abstract

A cell track contains not only a set of movements but also an order. We asked whether that order reveals behaviour that speed, endpoint displacement, and conventional persistence measures miss. For each trajectory, we compared the recorded sequence with an exact reference in which the same displacement steps were rearranged; the observed-minus-reference difference quantifies persistence created by step order. In 48,134 MCF10DCIS.com trajectories, most of a reproducible MYO10–collagen interaction in directional persistence disappeared when temporal order was removed. In 131 single MCF10A haptotaxis tracks, widening the fibronectin pattern did not produce a resolved global increase in order; it shifted short-timescale organization from the gradient axis toward lateral movement, with an intrinsic lateral-order difference of 0.2176 (95% hierarchical interval 0.1274–0.3171). In MDA-MB-231 cells migrating toward EGF, PFKL-N702T produced fewer up-gradient steps and shorter up-gradient runs while the mean-speed difference remained unresolved. These directional effects remained after comparisons balanced for measured speed and duration, whereas three cue-independent serial-order measures showed no reduction larger than a locked 0.10-SD margin at the track level. Across independent endothelial, epithelial, and cancer-cell archives, the same calculation distinguished sustained, short-lived, and sign-reversing regimes; a stationary angular hidden-state model failed all eight frozen long-lag targets. Temporal order therefore separates three biological questions: how much a cell moves, whether it continues moving persistently, and whether that persistence remains aligned with an environmental cue. We provide a documented workflow that calculates these readouts from standard x–y tracking tables.

**Impact statement:** The order of otherwise similar migration steps reveals whether cells merely continue moving or sustain commitment to an environmental cue.

## INTRODUCTION

Directed cell migration requires cells to detect environmental information, transmit it to polarity and cytoskeletal systems, and repeatedly convert that information into asymmetric movement (SenGupta et al., 2021). Speed, path length, endpoint displacement, mean-squared displacement (MSD), turning angles, and path-persistence measures remain indispensable (Gorelik and Gautreau, 2014; Maiuri et al., 2015).

However, each compresses a track and can discard the sequence in which individual movements occurred. Two cells can use the same set of step vectors, move at the same mean speed, and finish at the same endpoint, yet arrange those steps differently in time. That temporal arrangement can determine whether a cell sustains a direction, switches between directions, or alternates between polarity states.

Figure 1 introduces the analysis in biological terms. A migration step is the displacement between two consecutive frames. The observed track is compared with an exact shuffled-order reference containing the same steps in a different temporal order. Because the reference retains all recorded step lengths and directions, track duration, net displacement, and static mean direction, the observed-minus-reference difference isolates the contribution of temporal arrangement. We call this sequence-dependent persistence; the equivalent mathematical label in the source tables is sequence excess. Table 1 defines every recurring biological and statistical term in plain language, while Appendix 1 contains the equations and exact identities.

**Figure 1.**
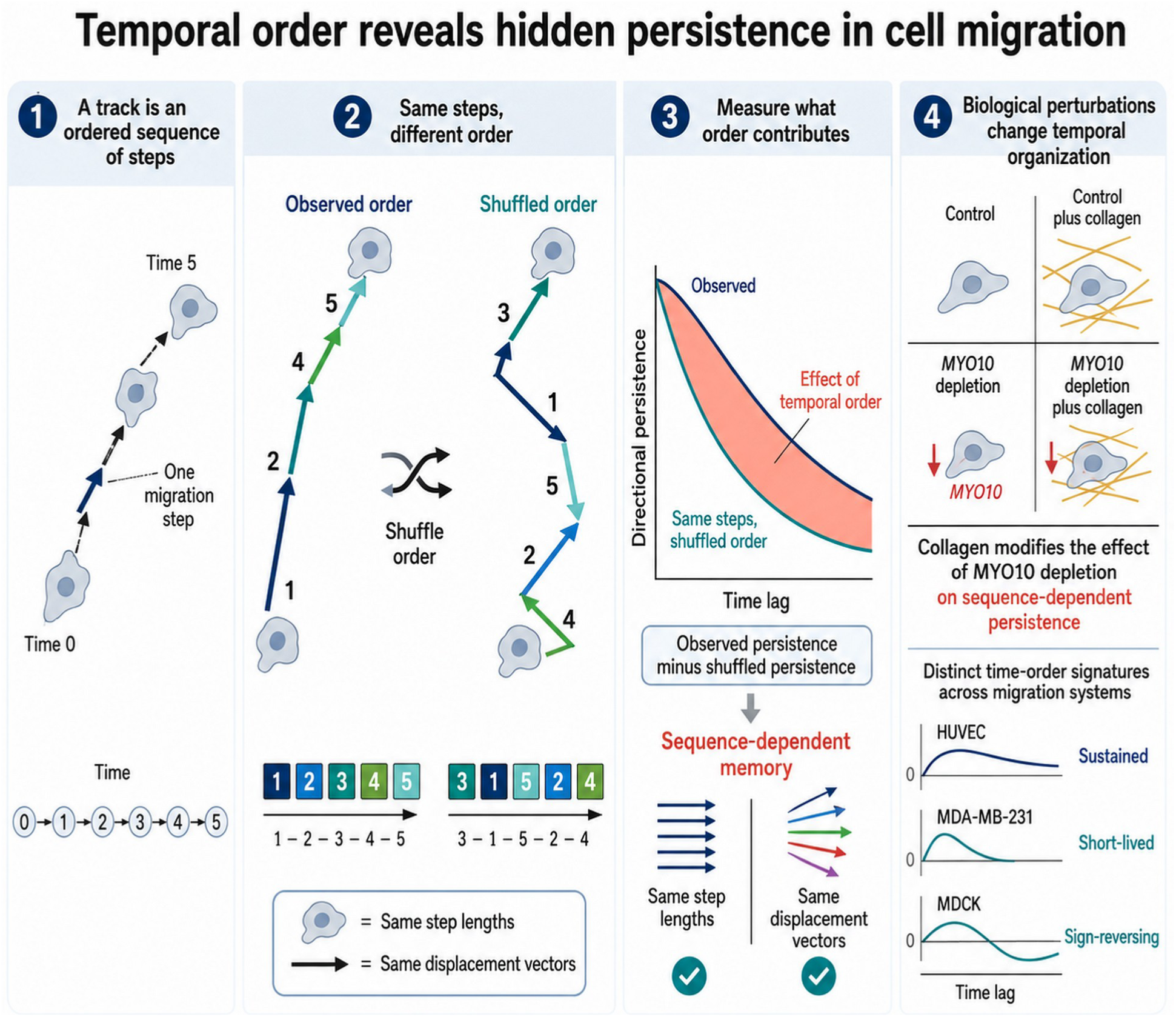
Plain-language overview of temporal-order analysis. (1) A track is represented as an ordered series of displacement steps. (2) The analytical order-null uses the same displacement vectors in a shuffled temporal order, preserving step lengths, directions, track length, static polarity, and endpoint displacement. (3) Observed persistence minus shuffled-order persistence is sequence-dependent persistence: the part of directional memory contributed by temporal arrangement. (4) Biological perturbations can alter temporal organization, and different migration systems can exhibit sustained, short-lived, or sign-reversing signatures.

**Table 1.** Plain-language glossary and statistical safeguards. Canonical terms are used consistently in the main text; mathematical symbols and derivations are given in Appendix 1.

| Term | Plain-language definition | What it distinguishes |
| --- | --- | --- |
| Track (trajectory) | The time-ordered list of a cell's recorded positions. | The complete recorded migration history of a single cell. |
| Migration step (increment) | The displacement vector between two consecutive imaging frames. | One movement, including both its length and direction. |
| Time lag | The time separating two steps whose similarity is being compared. | Short-timescale from long-timescale organization. |
| Directional persistence / VACF | The average similarity of displacement vectors separated by a chosen time lag; VACF is the velocity-autocorrelation form used here. | Continued movement in a similar direction from turning or reversal. |
| Static polarity | A track's overall mean displacement direction, irrespective of when individual steps occurred. | A persistent average direction from genuinely ordered sequences of steps. |
| Order-null (shuffled-order reference) | The exact expected persistence after uniformly rearranging the temporal positions of a track's recorded steps. | Step content and static polarity from effects caused by temporal arrangement. |
| Sequence excess / sequence-dependent persistence | Observed persistence minus the order-null value. These are the mathematical and biological names, respectively, for the same observed-minus-null contrast. | Persistence created specifically by the recorded order of steps. |
| Generic serial order (serial-order memory) | Sequence-dependent continuation without requiring movement toward a named cue. | General temporal persistence from cue-directed commitment. |
| Cue axis | The experimentally defined direction of a gradient or other directional signal. | Productive movement relative to the cue from movement in other directions. |
| Forward migration index (FMI) | Net displacement along a specified axis divided by total path length. | How efficiently movement is converted into progress along the cue axis. |
| Cue-aligned commitment / up-gradient run | A sustained sequence of steps directed toward the cue; a run is the consecutive block used to quantify it. | Persistent movement that remains productively aligned from generic persistence. |
| Axis contribution | Axis-specific sequence excess normalized by total movement activity; it combines how much movement occurs on that axis with how ordered it is. | The spatial allocation of temporal order among axes. |
| Intrinsic axis order | Axis-specific sequence excess normalized by activity on that same axis. | Order within the axis from the mere frequency of movement on that axis. |
| Accumulated displacement memory | The cross-step contribution through which correlations among earlier movements add to displacement over a longer window. | Immediate step activity from temporal correlations that accumulate over time. |
| Field of view (FOV) | One imaged region containing a group of tracked cells. | Cells sharing one acquisition context from cells in other images or repeats. |
| Hierarchical aggregation | Averaging at the levels set by the experimental design, such as track to FOV to biological repeat. | Biological replication from the potentially much larger number of tracked cells. |
| Factorial interaction | In a two-by-two experiment, the combined-condition response minus the no-interaction expectation from the two individual perturbations. | An additive or multiplicative no-interaction response from buffering/antagonism or enhancement. Here, a positive interaction under two suppressive main effects denotes buffering, not synergy. |
| Simultaneous confidence band | An uncertainty band constructed to cover an entire predeclared lag curve rather than one lag at a time. | A robust lag interval from isolated pointwise fluctuations. |
| HC3 robust interval | A regression confidence interval using a heteroskedasticity-consistent variance estimator that is less sensitive to unequal residual variance. | A covariate-adjusted association from an unadjusted group difference. |
| Propensity matching / overlap weighting | Two balance methods that compare groups with similar measured covariates: matching pairs comparable tracks, whereas overlap weighting emphasizes their common covariate range. | A directional phenotype from imbalance in measured speed and duration. |
| Non-inferiority margin | A predefined maximum reduction considered small enough to rule out a biologically meaningful loss. | Evidence of no meaningful reduction from a merely nonsignificant difference test. |

This distinction motivates a biological question rather than only a statistical one: do perturbations alter the ability to move, the tendency to continue in any direction, or the ability to keep persistent movement aligned with a specific cue? We use generic serial order for persistence that does not require a named cue and cue-aligned commitment for sustained sequences directed toward an extracellular signal. This separation is relevant because cue sensing and motility can be experimentally dissociated. Chemotactic systems can retain movement while losing directional sensing, and physical confinement can redirect the axes along which persistent movement is expressed (Skoge et al., 2014; SenGupta et al., 2021; Fortunato et al., 2025; Hansen and Webb, 2025).

Our discovery setting was a public four-condition MCF10DCIS.com wound-healing experiment combining MYO10 depletion and collagen exposure. MYO10 is an actin-based filopodial motor that supports integrin activation, lamellipodin recruitment, protrusion stability, and extracellular-matrix organization (Bohil et al., 2006; Miihkinen et al., 2021; Peuhu et al., 2022; Popović et al., 2023). Collagen simultaneously changes adhesive, geometric, and mechanical guidance (Paszek et al., 2005; Provenzano et al., 2008; Wolf et al., 2013; Licup et al., 2015). A two-by-two design allows a factorial interaction to ask whether the combined perturbation differs from the response expected by adding, or multiplying on a log scale, the two individual effects. The original manuscript identified a reproducible MYO10–collagen interaction but did not make sufficiently clear what the order-sensitive analysis added biologically.

We therefore extended the study to two direct tests of cue-guided single-cell behaviour. First, MCF10A cells migrating on fibronectin gradients of different widths provide a haptotaxis system in which confinement changes whether cells move longitudinally, oscillate, turn, or explore laterally (Fortunato et al., 2025). This asks whether geometry changes the amount of temporal order or redistributes it between the gradient and transverse axes. Second, MDA-MB-231 breast-cancer cells migrating toward an EGF gradient provide a chemotaxis system in which PFKL depletion, catalytic inactivation, pharmacological inhibition, and disruption of PFKL filament formation alter directional sensing. The PFKL-N702T mutant is especially informative because the source study reported impaired directional sensing without a loss of migration velocity (Hansen and Webb, 2025). This asks whether a cell can remain serially persistent yet fail to align that persistence with a chemoattractant.

Trajectory reanalysis also requires safeguards. Tracks are nested within image fields and biological repeats, identifiers may reset between movies, and the set of tracks available can shrink at long time lags. Treating all cells as independent biological replicates inflates precision. We therefore preserve the source hierarchy wherever it is recoverable; use lag-invariant cohorts, meaning the same eligible tracks are compared across the stated lag range; calculate analytical rather than Monte Carlo order-null values; test exact decomposition closure; and state explicitly when public files do not retain biological-replicate labels. A user-facing command-line workflow accepts x-y track tables and returns conventional metrics, order-null persistence, sequence excess, axis-resolved order, and cue-directed run statistics together with machine-readable audit files.

Here we show that temporal order provides a common but biologically interpretable coordinate across matrix, cytoskeletal, and metabolic perturbations. In the MYO10–collagen discovery experiment, step order carries most of a buffering interaction in directional persistence. In single-cell haptotaxis, geometry redistributes order between the gradient and lateral axes. In cancer-cell chemotaxis, PFKL filament disruption weakens cue-aligned runs without a resolved speed deficit or global loss of serial order. The same framework therefore distinguishes how much cells move, whether they persist, and whether that persistence remains committed to a cue.

### Reader guide: three questions asked of every track

**1. Motility.** How fast and how far does the cell move? Conventional outputs include speed, path length, displacement, and MSD.
**2. Generic persistence.** Does the recorded order keep successive movements aligned more than expected from the same steps rearranged in time? Sequence-dependent persistence answers this question without requiring a named cue.
**3. Cue-aligned commitment.** When a cue axis is known, does persistent movement remain directed toward it? FMI, up-gradient step fraction, and up-gradient run length answer this question.

**How a cell biologist can use the workflow**

**Input**: A standard delimited table containing track identifier, time, x, and y; optional columns can specify movie/FOV, condition, biological repeat, and cue direction.

**Output**: The workflow returns conventional migration measures, the exact shuffled-order reference, sequence-dependent persistence, axis-resolved order, and cue-directed run statistics, plus audit files that flag identifier reuse, missing frames, and unit inconsistencies.

**Inference**: Cell-level summaries can be calculated for any track table, but repeat- or FOV-level inference is enabled only when those experimental labels are supplied. The software does not manufacture biological replication from pooled cells.

## RESULTS

### Temporal order measures persistence contributed by sequence rather than step content

For each trajectory, we calculated conventional migration measures and a lag-resolved directional persistence curve. We then calculated the exact expectation after uniformly permuting the temporal positions of the recorded steps. No simulated track was required: the expectation follows analytically from the track’s mean step vector and mean squared step length. The order-null therefore retains whether a track has an overall preferred direction but removes whether similarly directed steps occurred in sustained blocks.

Sequence-dependent persistence is the observed curve minus this order-null curve. A positive value means that similarly directed movements were closer together in time than expected from the track’s static polarity alone. A negative value means that the observed order promoted alternation or reversal. Because long-window displacement is the sum of individual steps, the same distinction can be propagated exactly into direct step activity, order-null cross-step memory, and sequence-dependent displacement memory. All identities were required to close to machine precision before biological interpretation.

This construction makes the biological comparison explicit. Conventional speed asks how quickly a cell moves. Path persistence asks how directly the full track connects its start and endpoint. Generic serial order asks whether movements remain temporally aligned in any direction. Axis-resolved order asks where this organization is expressed. Cue-directed run statistics ask whether it is aligned with an experimentally defined signal.

### Corrected trajectory identities reveal a reproducible MYO10–collagen interaction

The discovery dataset contained field-local TRACK_ID values that reset between movies. Grouping only by condition and TRACK_ID merged unrelated cells: 4,581 identifiers were reused across fields of view (FOVs). Reconstruction with the supplied Unique_ID yielded 49,268 raw trajectory identities across 117 FOVs. After frozen continuity and minimum-length filtering, 48,134 identities remained: 14,219 shCTRL, 9,608 shCTRL+collagen, 14,331 shMYO10, and 9,976 shMYO10+collagen. The primary cohort used the longest retained contiguous segment from each identity (Figure 2A,B; Table 2).

**Figure 2.**
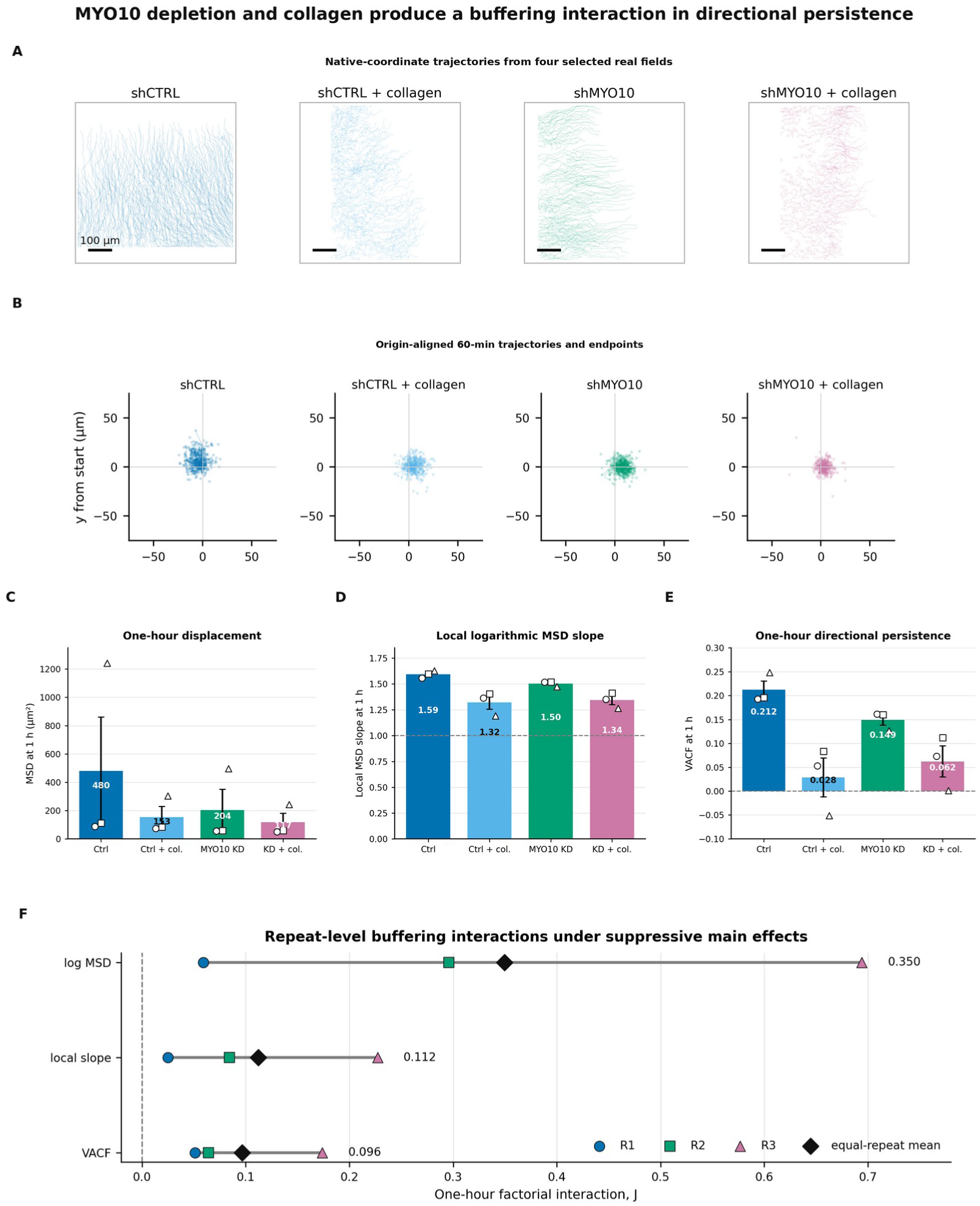
MYO10 depletion and collagen produce a buffering interaction in directional persistence. (A) Native- coordinate trajectories from one outcome-independently selected real FOV per condition. Coordinates and scale bars are data-derived; track lines were thinned only for legibility. (B) Origin-aligned trajectories over the first 60 min and all observed 60-min endpoints. (C-E) Equal-repeat condition means for one-hour MSD, the local logarithmic MSD slope, and normalized VACF; error bars are the standard errors across R1-R3 and open symbols show the three repeat estimates. The dashed lines mark a local slope of 1 in D and zero VACF in E. (F) Repeat-level factorial interactions for log MSD, local slope, and VACF. Colored symbols are R1-R3, horizontal lines span the repeat estimates, and black diamonds are equal-repeat means. Positive interaction under two suppressive main effects denotes buffering or antagonism on the analyzed scale, not positive synergy.

**Table 2.** Corrected primary cohort and one-hour observables. Values are repeat-equal-weighted condition means. α(1 h) is the local logarithmic MSD slope, not a global fitted exponent.

| Condition | Primary trajectory identities | FOVs per repeat | MSD at 1 h ($\mu\text{m}^2$) | Local $\alpha(1\text{ h})$ | VACF at 1 h |
| --- | --- | --- | --- | --- | --- |
| shCTRL | 14,219 | 9, 9, 9 | 480.004 | 1.5937 | 0.2123 |
| shCTRL + collagen | 9,608 | 10, 10, 10 | 152.929 | 1.3210 | 0.0285 |
| shMYO10 | 14,331 | 10, 10, 10 | 203.603 | 1.5040 | 0.1495 |
| shMYO10 + collagen | 9,976 | 10, 10, 10 | 116.951 | 1.3435 | 0.0621 |

Correct reconstruction changed the physical interpretation. At 60 min, local logarithmic MSD slopes were 1.594, 1.321, 1.504, and 1.343 for shCTRL, shCTRL+collagen, shMYO10, and shMYO10+collagen, respectively; all were superdiffusive rather than confined. Corresponding MSD values were 480.0, 152.9, 203.6, and 117.0 µm². Normalized velocity-autocorrelation values (VACF; the lagged similarity of displacement vectors after normalization by step activity) were 0.212, 0.028, 0.149, and 0.062 (Figure 2C- E; Table 2).

Collagen exposure and MYO10 depletion each suppressed displacement and persistence relative to shCTRL. Their combination nevertheless produced a positive factorial interaction: the combined condition was less suppressive than expected from the two individual perturbations on the analyzed scale. The repeat-level log-MSD interactions were 0.059, 0.296, and 0.694 (mean 0.350); exponentiation corresponds to approximately 1.42-fold the multiplicative no-interaction expectation. Mean interactions in the local MSD slope and normalized velocity autocorrelation were 0.112 and 0.096 (Figure 2F). We therefore use buffering interaction rather than synergy: both main effects are suppressive, but the combined suppression is weaker than expected.

Trajectory counts were strongly imbalanced among repeats, particularly because R3 contained many more tracks. All principal estimates therefore gave equal weight first to FOVs and then to the three source-labelled repeats. In a lag-invariant cohort, in which the same eligible trajectories supported every stated lag, the directional-persistence interaction was simultaneously robust from 30 to 150 min and positive in R1, R2, and R3 at 60 min (Figure 3A,F; Appendix 1—figure 4). All 117 leave-one-FOV-out calculations retained a positive mean. Matched early-versus-late windows did not support progressive strengthening during the movies; the interaction persisted across lag without a resolved temporal increase (Appendix 1—figure 5).

**Figure 3.**
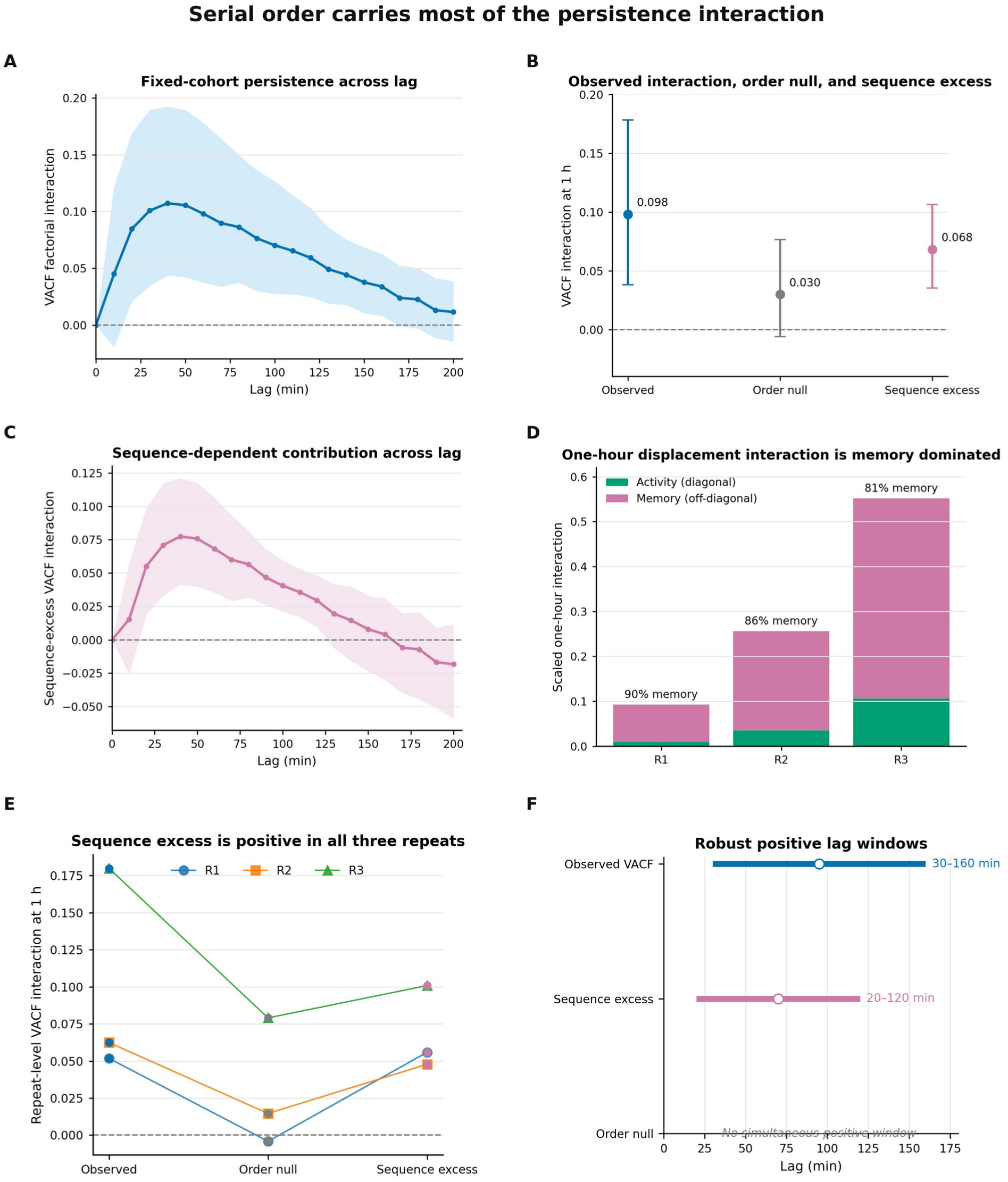
Serial order carries most of the persistence interaction. (A) Fixed-cohort VACF factorial interaction across lag with its pointwise 95% interval. (B) One-hour observed VACF interaction, analytical order-null interaction, and sequence-excess interaction with pointwise 95% intervals. (C) Sequence-excess interaction across lag with its pointwise 95% interval. (D) Exact one-hour displacement interaction decomposed into diagonal activity and off-diagonal memory for R1-R3; labels give the memory share of the direct interaction. (E) Repeat-level observed, order- null, and sequence-excess interactions at one hour. (F) Contiguous lag windows whose simultaneous bands remained above zero; no such window was detected for the order-null interaction.

### The MYO10–collagen interaction is carried mainly by the order of directions

At 60 min, the observed normalized persistence interaction was 0.0980. The order-null interaction was 0.0298 and had no simultaneously robust lag window. Sequence-dependent persistence was 0.0682, with a simultaneous 95% interval of 0.0209–0.1156 and the same positive sign in all three repeats (Figure 3B,E). The sequence-dependent component remained robust over 20–120 min, whereas the static order-null component did not (Figure 3C,F). Descriptively, temporal order accounted for approximately 70% of the observed 60-min interaction. This fraction is not an independent test, but it shows what the method adds beyond static polarity.

The displacement identity provided a second view. At 60 min, cross-step memory carried 90.1%, 86.4%, and 80.8% of the positive raw additive displacement interaction in R1–R3 (Figure 3D). Step separations of 20–120 min contributed positively to sequence-dependent memory accumulated over 120–200-min displacement windows (mean 0.158, bootstrap interval 0.074–0.242; Appendix 1—figure 6). Opportunity normalization showed that these separations contributed reproducibly but were not uniquely enriched relative to all valid separations. Thus, the data support intermediate-timescale organization without implying a sharply tuned molecular clock.

An exact speed–direction decomposition identified the robust carrier. The mean-speed directional term was simultaneously positive over 50–150 min, whereas speed-memory modulation and speed–direction coupling had no robust interval (Table 3; Appendix 1—table 3). This does not make speed biologically irrelevant; it means that among the algebraic components of lagged covariance, direction was the only component with repeat-consistent temporal support.

**Table 3.** Frozen evidence hierarchy. “Supported” denotes repeat-sign consistency, leave-one-repeat-out sign preservation, and simultaneous-band exclusion of zero where applicable.

| Claim | Status | Frozen interval or estimate | Interpretation |
| --- | --- | --- | --- |
| Superdiffusive corrected local slopes | Supported | $\alpha(1\text{ h})=1.32\text{--}1.59$ | Earlier confinement arose from cross-FOV identity merging |
| VACF factorial interaction | Supported | 30–150 min; $J(1\text{ h})=0.098$ | Buffering under suppressive main effects |
| Sequence order beyond static polarity | Supported | VACF 20–120 min; covariance 40–100 min | Increment order is load-bearing |
| Sequence-memory accumulation | Supported | 120–200 min | Serial correlations accumulate at long displacement lags |
| 20–120-min source contribution | Supported | 0.158; 95% interval 0.074–0.242 | Positive contribution to 120–200-min memory |
| Kernel-normalized source positivity | Supported | $8.63\times 10^{-4}$; $3.92\times 10^{-4}$–$1.34\times 10^{-3}$ | Positive contribution per kernel opportunity |
| Preferential source-band enrichment | Not supported | Difference interval crosses zero; signs +, +, – | No uniquely tuned 20–120-min scale |
| Early-to-late strengthening | Not supported | No robust gain window | Do not claim adaptation during imaging |
| Stationary-local generative explanation | Not supported | Linear 2/9; angular 0/8 targets passed | Signs/coarse memory do not reproduce quantitative hierarchy |
| Focal-cell self-inclusion | Supported | LOO total unchanged; shared-flow allocation $\Delta=1.4\times 10^{-5}$ at 1 h | Inclusive reference field does not generate the collective result |
| External framework transportability | Supported | 65 movies; robust horizons 60–120 min depending system | Framework distinguishes sustained, short, and sign-reversing regimes |

The directional interaction was present at both collective and cell-relative scales. A leave-one-cell-out field velocity was calculated from all other contemporaneous cells in the same FOV, preventing the focal cell from contributing to its own reference flow. Shared FOV flow and cell-relative motion contributed positively, whereas their summed cross term was negative over much of the analyzed lag range (Appendix 1—figure 7A-C). Removing focal-cell self-inclusion changed the 60-min allocation by less than 4.1×10⁻⁴ and left total persistence unchanged. The negative cross term is a kinematic covariance, not evidence of biochemical compensation.

The interaction also survived matching and exclusion controls for measured FOV occupancy, nearest-neighbour spacing, image-edge distance, and trajectory duration. Across these analyses, the 60-min interaction remained positive in every repeat and ranged from 0.087 to 0.103, compared with 0.098 in the primary analysis (Table 3; Appendix 1—table 5). Unmeasured matrix architecture or cell state can still contribute, but the signal is not explained by the measured context variables.

### Physical confinement redistributes temporal order during single-cell haptotaxis

To test whether temporal order maps onto an intuitive single-cell behaviour, we analyzed 131 complete MCF10A trajectories from fibronectin gradients confined to widths of 20, 40, 60, 80, or 250 µm (Fortunato et al., 2025). Sequence-dependent persistence was positive at every width, indicating that temporal order survived substantial changes in accessible geometry. The central contrast was not a monotonic gain or loss of total order. Instead, width changed the axis along which order was expressed (Figure 4A-D; Table 4).

**Figure 4.**
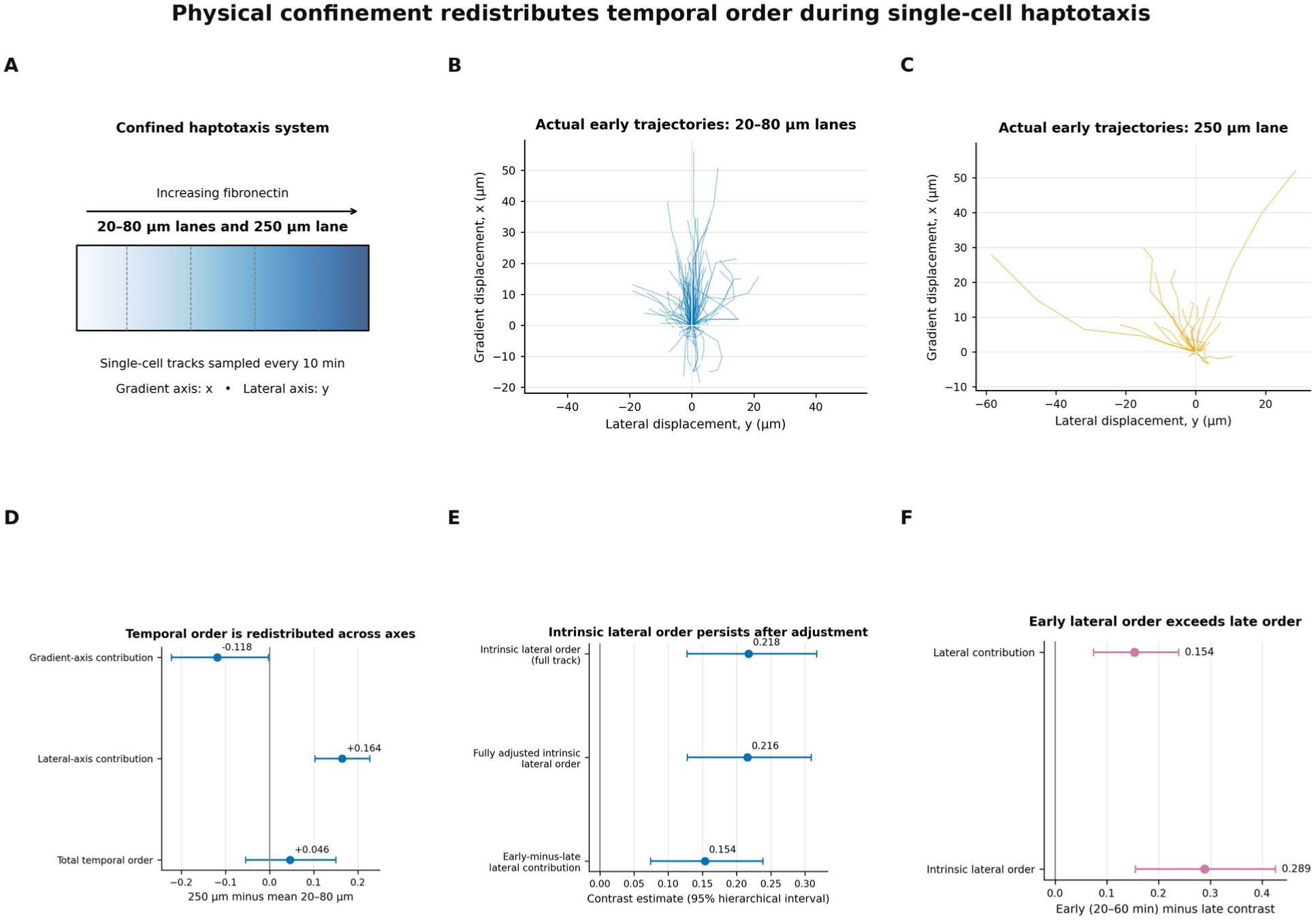
Physical confinement redistributes temporal order during single-cell haptotaxis. (A) Experimental geometry: cells migrate on confined fibronectin-gradient patterns spanning 20–80 µm lanes and a 250-µm lane. (B,C) Actual 20–60-min trajectories from the 20–80-µm and 250-µm geometries, respectively, origin-aligned with the gradient plotted vertically and the lateral axis horizontally. (D) Wide-minus-confined contrasts for gradient-axis contribution, lateral-axis contribution, and total sequence-dependent order at 20–60 min. (E) Primary intrinsic lateral order, its fully adjusted estimate, and the early-minus-late lateral-contribution contrast. (F) Direct early-minus-late contrasts for lateral contribution and intrinsic lateral order. Points are estimates and bars are hierarchical 95% intervals. Panel A is schematic; panels B–F use deposited coordinates and validated analysis tables.

**Table 4.** Locked single-cell haptotaxis contrasts. Positive values denote greater order in the 250-µm geometry than the mean of 20–80-µm lanes unless stated otherwise. Intervals are hierarchical 95% intervals.

| Endpoint | Contrast | Estimate | 95% interval | Interpretation |
| --- | --- | --- | --- | --- |
| Lateral axis contribution, 20–60 min | 250 $\mu\text{m}$ – mean(20–80 $\mu\text{m}$) | +0.1643 | 0.1025 to 0.2271 | More short-lag lateral allocation of order in wide geometry |
| Gradient-axis contribution, 20–60 min | 250 $\mu\text{m}$ – mean(20–80 $\mu\text{m}$) | –0.1182 | –0.2222 to –0.0021 | Less short-lag gradient-axis allocation of order |
| Total order, 20–60 min | 250 $\mu\text{m}$ – mean(20–80 $\mu\text{m}$) | +0.0461 | –0.0542 to 0.1502 | No resolved global increase |
| Intrinsic lateral order, full track | 250 $\mu\text{m}$ – mean(20–80 $\mu\text{m}$) | +0.2176 | 0.1274 to 0.3171 | Lateral steps themselves are more ordered |
| Fully adjusted intrinsic lateral order | 250 $\mu\text{m}$ – mean(20–80 $\mu\text{m}$) | +0.2159 | 0.1277 to 0.3092 | Not explained by measured activity, speed, persistence, or duration |
| Lateral early-minus-late contrast | (20–60 min) – (60–180 min) | +0.1538 | 0.0745 to 0.2386 | Redistribution is stronger at short lags |
| Observation-horizon contrast | 6 h – 3 h | Unresolved | Interval included zero | No evidence for a discrete six-hour onset |

At short lags of 20–60 min, the 250-µm geometry had a greater lateral-axis contribution than the mean of the 20–80-µm lanes (difference 0.1643, 95% hierarchical interval 0.1025–0.2271). The gradient-axis contribution shifted in the opposite direction (−0.1182, interval −0.2222 to −0.0021), while the total-order contrast was unresolved (0.0461, interval −0.0542 to 0.1502; Figure 4D). Because an axis contribution combines movement allocation with intrinsic order, this result shows that wide geometry reallocates short-timescale temporal organization from the gradient axis into lateral exploration; the next analysis tests whether lateral movements themselves are also more ordered.

The lateral effect was not only a consequence of wider cells taking more transverse steps. After normalizing by lateral activity, the intrinsic lateral-order contrast between 250 µm and the mean of 20–80 µm was 0.2176 (95% interval 0.1274–0.3171). Adjustment for lateral activity, speed, path persistence, and duration gave 0.2159 (0.1277–0.3092). The direct early-minus-late contrast was 0.1538 (0.0745–0.2386), supporting concentration of the lateral redistribution at shorter lags (Figure 4E,F; Appendix 1—figure 11). A separate 6- h-minus-3-h comparison remained unresolved, so the data do not establish a discrete onset at six hours.

Directional reversals provided a second visible behaviour. In the finalized analysis, reversal-aligned trajectories were compared using a common complete-case cohort and matched pseudo-event times from the same tracks. The true reversal window showed a transient reorganization of step order beyond the pseudo-event reference. This should not be interpreted as a simple increase in one-direction persistence: outgoing and incoming polarity states become temporally segregated during the switch. Together, the geometry and reversal results show where and when temporal order becomes biologically expressed in single cells.

### PFKL filamentation supports cue-aligned commitment without reducing generic serial order

We next analyzed MDA-MB-231 breast-cancer cells migrating for 16 h toward a negative-x EGF gradient in a public PFKL study (Hansen and Webb, 2025). The deposited tracking CSVs contained 1,691 reconstructed tracks across acute and stable PFKL depletion, catalytic-dead PFKL-H199Y, PFK15 treatment, and filament-incompetent PFKL-N702T. The source article reports three biological replicates per condition, but replicate identity was not retained in the compiled trajectory labels. Accordingly, the intervals below quantify track-level uncertainty and the five perturbation series are treated as mechanistically distinct within-study comparisons, not five independent studies.

The primary comparison contained 150 PFKL-WT and 150 PFKL-N702T tracks. The mean-speed difference was unresolved (N702T−WT, −0.677 µm h⁻¹; track-bootstrap 95% interval −1.424 to 0.121). In contrast, the EGF-axis forward migration index (FMI; net cue-axis displacement divided by total path length) fell by 0.278 (−0.358 to −0.200), the up-gradient step fraction by 0.166 (−0.210 to −0.121), and mean up-gradient run length by 0.666 steps (−1.127 to −0.204). Final cue-directed displacement fell by 24.31 µm (−30.53 to −17.81). Thus, N702T cells remained motile but were less able to sustain movement in the productive cue direction (Figure 5B; Table 5).

**Figure 5.**
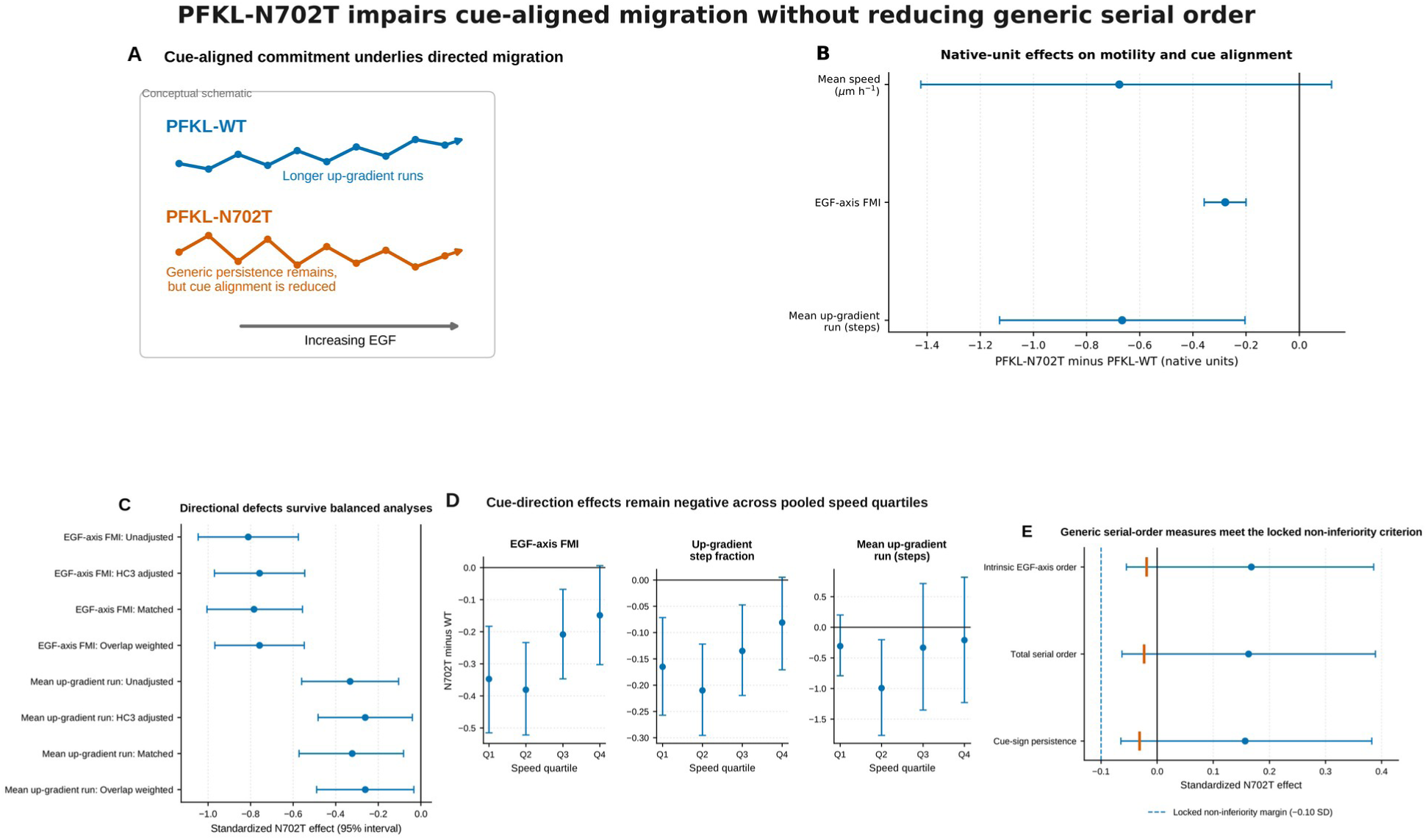
PFKL-N702T impairs cue-aligned migration without reducing generic serial order. (A) Conceptual distinction between persistent movement aligned with the EGF gradient and persistent movement that is less productively aligned. (B) Track-level N702T-minus-WT estimates for mean speed, EGF-axis FMI, and mean up-gradient run in native units. (C) Directional effects in unadjusted, HC3-adjusted, propensity-matched, and overlap-weighted analyses. (D) Cue FMI, up-gradient step fraction, and mean up-gradient run effects across pooled speed quartiles; point estimates are negative in every quartile, while some quartile-specific intervals cross zero. (E) Standardized effects for generic serial-order measures. The dashed line is the locked −0.10-SD non-inferiority margin and orange ticks mark one-sided 95% lower bounds. Biological-replicate identity was unavailable in the public trajectory labels, so intervals are track-level. Panel A is schematic; panels B–E are plotted from the validated Stage 3 tables.

**Table 5.**
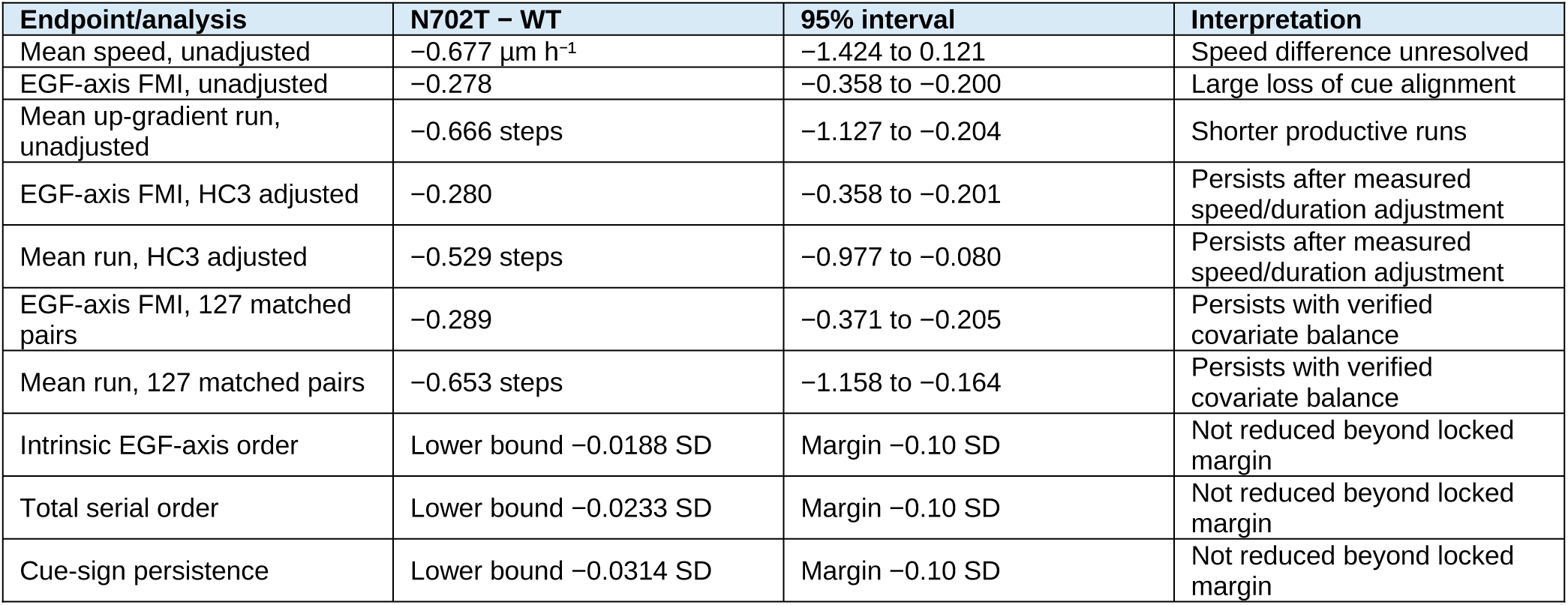
Primary PFKL-N702T versus PFKL-WT chemotaxis results. Unadjusted intervals are track- bootstrap intervals; adjusted intervals are HC3, paired-bootstrap, or weighted HC3 intervals. Biological- replicate identity was unavailable.

A locked post hoc robustness analysis was fixed before the final execution after the preceding exploratory audit. HC3 regression, which provides a covariate-adjusted interval robust to unequal residual variance, included N702T status, log mean speed, and duration but not the nearly collinear number of positions. The cue-FMI coefficient was −0.280 (95% interval −0.358 to −0.201) and the mean-run coefficient was −0.529 steps (−0.977 to −0.080). Propensity matching paired tracks with similar measured speed and duration and retained 127 pairs with a maximum absolute standardized covariate difference of 0.0089; matched effects were −0.289 for cue FMI and −0.653 steps for mean run. Overlap weighting emphasized the shared covariate range and achieved near-exact balance, giving corresponding effects of −0.280 and −0.529 steps (Figure 5C; Appendix 1—figure 12A,B). Across pooled speed quartiles, every point estimate for cue FMI, up-gradient step fraction, and mean up-gradient run was negative, although several quartile-specific intervals crossed zero; this supports directional consistency rather than significance in every quartile (Figure 5D). These analyses show that the directional phenotype is not explained by measured speed and duration differences; they do not establish a causal pathway independent of speed.

Generic serial organization did not fall with N702T. Intrinsic EGF-axis order, total serial order, and cue-sign persistence had positive point estimates. Rather than treating a nonsignificant difference as proof of preservation, we used a non-inferiority test: a reduction larger than 0.10 standard deviations was defined as the smallest meaningful loss for the final locked analysis. For all three measures, the one-sided 95% lower confidence bound exceeded the −0.10-SD margin (−0.0188, −0.0233, and −0.0314 SD, respectively; Figure 5E; Appendix 1—figure 12E). We therefore conclude at the track level that none of these generic serial-order measures was reduced by 0.10 SD or more. The distinction is not ordered versus disordered migration: N702T cells retained generic temporal continuation but aligned it less efficiently with the chemoattractant.

The same directional pattern appeared across all five PFKL perturbation series. Confidence intervals for cue FMI and up-gradient step fraction were negative for acute depletion, stable depletion, catalytic inactivation, pharmacological inhibition, and N702T (Appendix 1—figure 12D). This directionally concordant within-study evidence links perturbations of PFKL abundance, catalytic function, pharmacological activity, and filament formation to impaired cue-directed migration. The speed-balanced N702T comparison provides the clearest demonstration of what temporal-order analysis adds.

### Serial-order signatures differ among endothelial, epithelial, and cancer migration systems

We retained the earlier frozen evaluation of public TrackMate movies to test whether long-lived positive sequence-dependent persistence was universal. The fixed cohort contained 26,417 trajectories from 17 MDA-MB-231 movies, 18,641 from 18 HUVEC exports, 147,710 from 15 MDCK-bulk movies, and 103,844 from 15 MDCK-edge movies. Exact 60-min decomposition closure was zero in every movie.

HUVEC trajectories showed sustained positive sequence-dependent persistence through 120 min. The independent MDA-MB-231 archive showed a weaker, shorter positive regime. MDCK bulk and edge regions were positive at early lags but became negative at longer lags, indicating temporal alternation rather than persistent continuation (Figure 6A-C; Table 7). At 60 min, mean sequence excess was 0.3240 for HUVEC, 0.00659 for MDA-MB-231, 0.0650 for MDCK bulk, and 0.0564 for MDCK edge; all simultaneous intervals excluded zero. Exact 60-min decomposition further separated order-null and serial-order displacement contributions across systems (Figure 6D; Appendix 1—figure 10).

**Figure 6.**
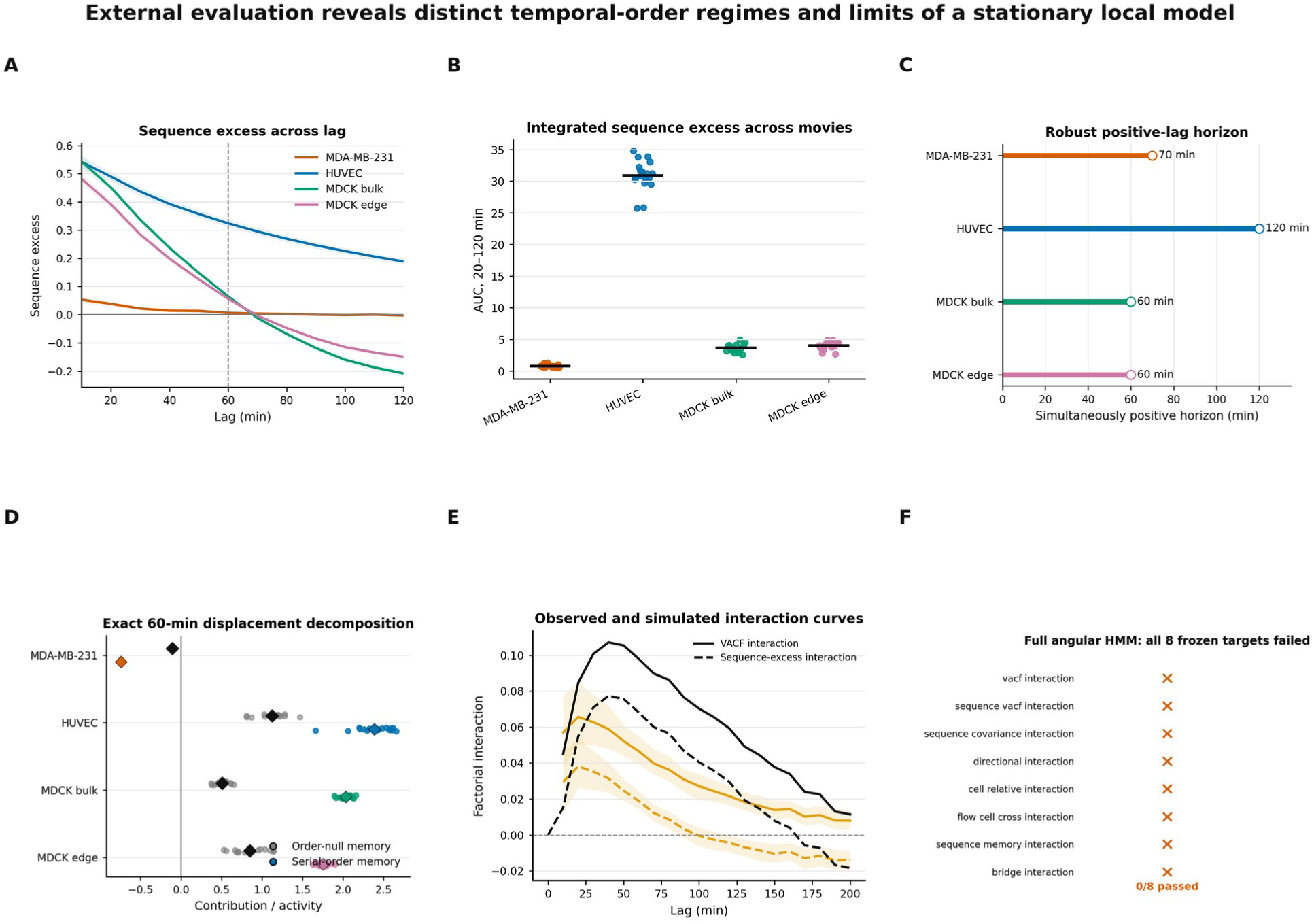
External evaluation reveals distinct temporal-order regimes and limits of a stationary local model. (A) System-level sequence-excess curves across lag with simultaneous 95% bands. (B) Movie-level sequence-excess AUC over 20–120 min; horizontal black lines denote system means. (C) Last lag whose simultaneous sequence- excess band remained above zero. (D) Movie-level order-null and serial-order contributions to exact 60-min displacement, normalized to activity; diamonds are system means. (E) Frozen observed VACF and sequence-excess interactions compared with predictive means and 95% intervals from the full angular hidden-state model. (F) Target-level validation for the full angular model: none of the eight frozen long-lag targets passed. Cross-system comparisons are descriptive because species, substrate, acquisition, and laboratory differ.

**Table 6.** Generative-model falsification summary. Predictive coverage, not sign frequency, was the primary criterion.

| Model | Fitting target | Predictive experiments | Frozen targets passed | Primary conclusion |
| --- | --- | --- | --- | --- |
| Linear active-memory | Local flow and residual transitions | 128 per scenario | 2/9 | Collective flow and integrated sequence memory reproduced; detailed hierarchy failed |
| Angular hidden-state | Local turns and state transitions; measured speed/flow conditioned | 128 per scenario | 0/8 | Two-state local angular likelihood improved, but all eight long-lag quantitative targets failed |

**Table 7.** Frozen external-validation summary. Sequence excess is observed normalized covariance minus the analytical within-trajectory order-null. Intervals are simultaneous 95% bands at 60 min. The serial-order contribution is the exact 60-min displacement-memory component normalized to activity; it integrates pair separations shorter than 60 min.

| System | Public XMLs | Fixed tracks | Sequence AUC 20–120 min | Sequence excess at 60 min [simultaneous 95%] | Robust positive horizon (min) | Serial-order contribution at 60 min |
| --- | --- | --- | --- | --- | --- | --- |
| MDA-MB-231 | 17 | 26,417 | 0.788 | 0.0066 [0.0023, 0.0109] | 70 | −0.741 |
| HUVEC | 18 | 18,641 | 30.917 | 0.3240 [0.3108, 0.3373] | 120 | 2.387 |
| MDCK bulk | 15 | 147,710 | 3.634 | 0.0650 [0.0595, 0.0705] | 60 | 2.037 |
| MDCK edge | 15 | 103,844 | 4.015 | 0.0564 [0.0508, 0.0620] | 60 | 1.755 |

These comparisons demonstrate transportability of the calculation, not a controlled cell-type experiment. Cell type, species, substrate, imaging, and tracking procedures differ together. Their value is therefore conceptual: a single statistic can distinguish sustained, short-lived, and sign-reversing temporal organization without assuming that one regime is universal.

### Stationary local models reproduce fragments but not the full empirical hierarchy

We finally asked whether local stationary generators could reproduce the full lag-resolved discovery hierarchy. A linear active-memory model fitted to one- and two-step transitions passed only two of nine frozen targets: shared-flow interaction and integrated sequence-dependent displacement memory. It failed the full persistence curve, sequence-dependent covariance, directional and cell-relative interactions, the negative cross term, and the exact pair-separation bridge. A no-coupling ablation reproduced the expected sign hierarchy more often than the full model, preventing a necessity claim for the fitted linear coupling (Table 6; Appendix 1—tables 6 and 7). Across the related fractional active-memory model zoo, posterior-predictive errors remained condition- and endpoint-dependent (Appendix 1—figure 8).

A nonlinear angular hidden-state model improved held-out local turning likelihood over a one-state model, but it failed all eight quantitative long-lag targets (Figure 6E,F; Appendix 1—figure 9). The result is not evidence for a uniquely non-Markovian molecular mechanism. It shows that matching local steps, signs, or integrated summaries is insufficient: a useful migration model must reproduce where temporal order is carried, how it accumulates, and how it is distributed between shared and cell-relative motion.

## DISCUSSION

The revised analysis answers a simple biological question: when two cells move with similar step content, what does the order of those steps reveal? Across the systems examined here, temporal order separates three quantities that are often conflated—motility, generic persistence, and commitment to an external cue. The MYO10–collagen experiment shows that a perturbation interaction can reside mainly in the sequence of directions. The haptotaxis experiment shows that geometry can redistribute where order is expressed without clearly increasing total order. The PFKL experiment shows that cells can remain motile and generically persistent while losing sustained alignment to a chemoattractant.

The MYO10–collagen result remains the discovery setting and has the strongest recovered experimental hierarchy. Equal-FOV and equal-repeat inference, fixed cohorts, leave-one-FOV-out analysis, and measured-context matching support a repeat-consistent buffering interaction. The order-null makes the additional contribution explicit: most of the 60-min interaction is absent when the same steps are temporally rearranged. Exact speed–direction and collective decompositions then locate the phenotype kinematically. Directional organization is robust; speed-memory terms are not. Shared flow and cell-relative motion both contribute, with a negative cross covariance. These are physical descriptions of the tracks, not direct measurements of molecular feedback.

MYO10-dependent filopodia provide a plausible route by which collagen context could influence this organization. MYO10 supports integrin activation, lamellipodin recruitment, protrusion stability, and basement-membrane organization, while collagen changes resistance and guidance. A cell that repeatedly samples and stabilizes protrusions may express these processes as blocks of similarly directed steps. The current tracks, however, contain no simultaneous adhesion, force, protrusion, or matrix-remodelling measurement. The study therefore identifies an order-sensitive MYO10–collagen phenotype and a testable hypothesis rather than a direct MYO10–collagen biochemical mechanism.

The haptotaxis analysis provides a more visible interpretation. Narrow lanes constrain the available directions, whereas wide patterns permit lateral turns and exploration. A resolved global change in total sequence-dependent order was not detected. Instead, its directional allocation changed: the short-lag gradient-axis contribution decreased while the lateral contribution increased. Intrinsic lateral normalization and multivariable adjustment showed that the finding was not merely the trivial result of taking more lateral steps. The transient reorganization around true reversals further connects the estimator to polarity-state transitions. Physical geometry therefore regulates where temporal persistence can be expressed.

The PFKL analysis provides a complementary biochemical example. The source study showed that PFKL activity and lamellipodial localization support directional migration and that N702T impairs directional sensing without reducing velocity. Our reanalysis resolves the temporal structure behind that statement. N702T cells made fewer up-gradient steps, sustained shorter productive runs, and ended with much less cue-directed displacement. These effects survived comparisons balanced for measured speed and duration, whereas generic serial-order measures met the locked track-level non-inferiority criterion. The pattern is consistent with compartmentalized glycolysis supporting the alignment of persistent movement with a cue rather than simply determining bulk motility; the trajectory data do not by themselves establish that molecular mechanism.

The PFKL inference has an important boundary. Biological-replicate labels are absent from the deposited tracking tables, so track-level intervals cannot replace replicate-level inference. The five perturbation series also arise from one source study. We use them as concordant mechanistic comparisons, not independent replications. The Stage 3 robustness plan was data-informed and then locked before final execution; it is not described as prospectively prespecified. Likewise, adjustment for speed supports the statement that measured speed differences do not explain the directional phenotype, not a causal claim that speed is irrelevant.

The cross-system results show why one persistence time is not always sufficient. Positive sequence-dependent persistence can remain sustained, decay rapidly, or become negative. A negative value is not ’less migration’; it means that the observed sequence favours alternation or reversal relative to the static track content. This language is intentionally descriptive. It avoids assuming a universal memory timescale or assigning every temporal signature to one molecular feedback loop.

The practical value of the method depends on adoption. The revised workflow therefore begins from standard tracking tables rather than microscopy-specific data structures. A minimal input contains track identifier, time, x, and y, with optional movie, condition, replicate, and cue-axis columns. The software returns a quality-control report, conventional migration metrics, observed and order-null persistence, sequence excess, axis-resolved order, and cue-run statistics. The analytical null avoids expensive permutation loops, and all output tables include input hashes, configuration, and random seeds. Exact factorial and hierarchical inference still require the experimental design to be specified correctly; the software cannot recover biological replication that is absent from source metadata.

Several limitations remain. The discovery experiment has only three highest-level repeats. The haptotaxis hierarchy partly relies on deposited block labels and a modest number of complete tracks. The PFKL public files do not preserve replicate identity. Cross-system comparisons confound biological and technical differences. The current analysis is kinematic: it does not measure signalling activity, force, adhesions, metabolic flux at the leading edge, or polarity dynamics in the same cells. Prospective experiments should combine 10-min-or-faster tracking with rescue perturbations and simultaneous reporters of protrusion, adhesion, traction, polarity, or localized ATP production.

In conclusion, temporal order is not an abstract replacement for established migration metrics. It is a complementary coordinate that asks whether recorded movements were arranged in a biologically productive sequence. Matrix context, physical geometry, and localized metabolism can affect different parts of that organization. By separating generic continuation from cue-aligned commitment, the framework provides a tangible readout for experiments in which speed and endpoint displacement alone cannot explain how cells navigate.

## MATERIALS AND METHODS

### Analysis overview and user-facing workflow

The common analysis treats a trajectory as an ordered sequence of two-dimensional displacement vectors. A documented command-line workflow accepts delimited tracking tables with track identifier, time, x, and y columns and optional movie, condition, biological-repeat, and cue-axis labels. The workflow audits identifier reuse, time continuity, missing frames, coordinate scale, and track duration; calculates conventional migration metrics; and writes observed persistence, analytical order-null persistence, sequence excess, axis-resolved order, and cue-directed run statistics. Design-aware modules perform FOV/repeat aggregation, factorial contrasts, hierarchical bootstrap, matching, or track-level robustness analyses only when the required metadata are available.

Every frozen run wrote source-data tables, input and output SHA-256 manifests, configuration, random seeds, software versions, numerical checks, and figure-rendering code. The order-null and displacement- memory identities were unit tested and required machine-precision closure. The accompanying peer-review package contains the user workflow, worked examples, deterministic tests, source tables, rendering code, and versioned result archives.

### Dataset, experimental design, and analysis unit

We analyzed public trajectory tables from the MYO10–collagen MCF10DCIS.com wound-healing experiment reported by Peuhu et al. (2022), compiled through CellTracksColab (Gómez-de-Mariscal et al., 2024), and deposited on Zenodo (Jacquemet et al., 2024). The source experiment followed shCTRL and shMYO10 MCF10DCIS.com cells with or without collagen overlay, acquired at 10-min intervals over 14 h, and organized into four conditions across three biological repeats as defined by the original investigators. The public record contained 117 FOVs. The present work is a secondary analysis of trajectory tables; no raw images were resegmented and no new experiments were performed. Original culture, treatment, imaging, and tracking procedures remain those of the primary publication and data record. The highest available inferential unit was the source-labelled biological repeat; trajectories were not treated as independent biological replicates. The operational independence of R1–R3 could not be verified beyond the public metadata, and the source quality-control analysis identified R3 as disproportionately large and phenotypically distinct.

The design was represented as a two-by-two factorial perturbation with MYO10 depletion M∈{0,1} and collagen exposure C∈{0,1}. All primary estimates were calculated within each repeat after equal-weight aggregation across FOVs. The final repeat estimate was the arithmetic mean of the three repeat-level values. This hierarchy prevents the largest repeat or the most densely tracked FOV from dominating an effect estimate.

### FOV-aware trajectory reconstruction and continuity audit

The source TRACK_ID resets between image fields. A track key formed only from condition and TRACK_ID therefore merges unrelated trajectories from different FOVs. We audited TRACK_ID reuse and reconstructed trajectories using the supplied Unique_ID, which encodes FOV/file identity together with the field-local track identifier. Rows were sorted by trajectory identity and time. Duplicate observations at identical times were audited, and trajectories were split at temporal discontinuities rather than joined across gaps.

After frozen continuity and minimum-length filters, a raw trajectory identity could yield one or two retained contiguous segments. Segment-level analyses used all retained segments where specified. The primary one-identity-one-contribution cohort selected the longest retained segment for each raw trajectory identity, with deterministic tie resolution. This yielded 48,134 trajectory identities and 3,123,278 observations. The staged reconstruction workflow is summarized in Appendix 1—table 1, and identity reconciliation, exclusion accounting, and condition-by-repeat counts are detailed in Appendix 1—table 2.

Trajectory visualization used the corrected physical coordinates without reconstruction of cell shapes or microscopy backgrounds. For Figure 2 and Appendix 1—figure 2, one FOV per condition was selected by an outcome-independent rule: among eligible FOVs, the FOV whose corrected track count was closest to the condition median was chosen, with deterministic tie resolution. All four selected FOVs happened to be source-labelled R2. Native-coordinate panels were deterministically thinned only to prevent overplotting; origin-aligned panels display a deterministic subset of track lines and all eligible 60-min endpoints. A strict filename-level match between these FOV identifiers and the available raw image inventory was not established, so no microscopy image was inserted.

### Drift-corrected positions and increments

For each retained contiguous segment, mean Cartesian velocity was calculated as endpoint displacement divided by segment duration. Within each repeat–condition–FOV unit, the drift vector was the componentwise median of those segment mean velocities. Let r(t) denote the centered trajectory position and t_₀_ its start time. The constant FOV-level translation accumulated from t_₀_ to t was subtracted to obtain the drift-corrected position:

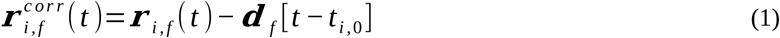

Migration increments at the nominal sampling interval Δt=10 min were

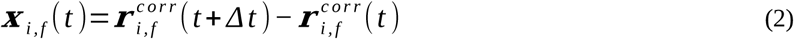

This correction removes only a time-independent FOV-level translational component; time-varying shared- field fluctuations remain available to the collective-flow decomposition. Raw absolute x and y coordinates were retained separately for nearest-neighbor, spatial-correlation, and image-edge analyses. Drift-corrected increments were used for the primary temporal observables; raw-versus-corrected sensitivity was evaluated before the evidence hierarchy was frozen.

### Mean-squared displacement, local logarithmic slope, and velocity autocorrelation

For a trajectory with N increments, the time-averaged squared displacement over n increments was computed from all valid origins:

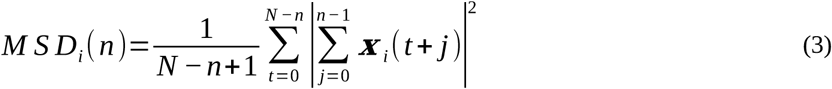

Trajectory values were averaged equally within FOV, FOVs equally within repeat-condition strata, and repeats equally for condition summaries. The local logarithmic MSD slope was not obtained from a single global power-law fit; it was evaluated numerically on each FOV-level curve:

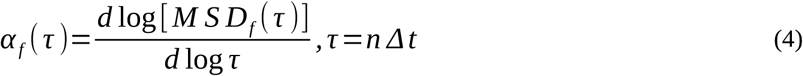

The normalized velocity autocorrelation at lag n was calculated using the trajectory-specific mean squared increment magnitude:

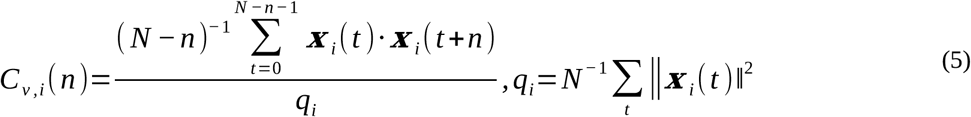

Additional descriptive observables included speed, radius of gyration, straightness, the non-Gaussian parameter, and ergodicity-breaking summaries. These were retained as diagnostics but were not promoted to primary claims when they failed the frozen repeat/sign/simultaneous-band criteria.

### Factorial interaction and scale interpretation

For any condition-level observable y, with indices 00=shCTRL, 01=shCTRL+collagen, 10=shMYO10, and 11=shMYO10+collagen, the factorial interaction was

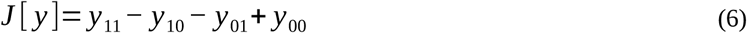

For MSD, the primary one-hour contrast was applied to log MSD, making the no-interaction model multiplicative:

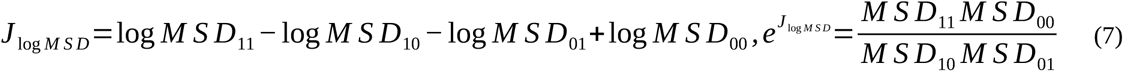

Because collagen exposure and MYO10 depletion both reduced displacement and persistence, J>0 denotes a combined response less suppressive than the additive or multiplicative no-interaction expectation on the relevant scale. We therefore use buffering or antagonism, not positive synergy.

### Hierarchical bootstrap, simultaneous bands, and robustness gates

The bootstrap resampled biological repeats with replacement and, within each selected repeat, resampled FOVs separately inside each condition. Pointwise 95% intervals were percentile intervals. For a lag curve J(τ), simultaneous bands used the bootstrap distribution of the maximum absolute standardized deviation across the predeclared lag grid (Efron and Tibshirani, 1993; Westfall and Young, 1993). Principal discovery stages used 20,000 replicates for one-hour factorial and analytical order-null inference and 10,000 replicates for lag-dependent and integrated decompositions, with seed 20260714. The leave-one-cell-out and external-validation analyses used 10,000 replicates with seed 20260716.

A contiguous lag interval was called robust only when (i) the interaction had the same sign in all three repeats, (ii) for each omitted repeat, the mean interaction of the two retained repeats had the same sign as the omitted repeat, and (iii) the simultaneous band excluded zero. Leave-one-FOV-out calculations removed each of the 117 FOVs in turn and recomputed the complete hierarchy. With three biological repeats, these sign, interval, and sensitivity criteria were emphasized over asymptotic p values.

### Lag-invariant cohorts and temporal-emergence test

Long-lag estimates can change composition as short tracks drop out. Fixed cohorts therefore required a minimum trajectory length before any lag-specific calculation. The primary full fixed cohort required at least 25 points for VACF and 24 points for displacement-memory analysis. Additional 36- and 48-point cohorts tested stricter censoring. For each raw trajectory identity, only the longest retained segment entered these cohorts.

Temporal emergence was tested by comparing early and late windows of equal length within the same trajectories. Repeat-level late-minus-early factorial contrasts were bootstrapped with the same hierarchy. This test addresses whether a broader late-lag robust interval reflects true strengthening during imaging rather than a stable interaction observed with different windows.

### Analytical temporal-order null

For a trajectory with N increments, define the mean increment vector mᵢ = N⁻¹Σₜxᵢ(t) and the mean squared increment magnitude qᵢ = N⁻¹Σₜ‖xᵢ(t)‖². Uniform random permutation preserves the complete increment multiset, trajectory length, net displacement, and static mean polarity while removing temporal order. The exact expected dot product between any two distinct permuted increments is

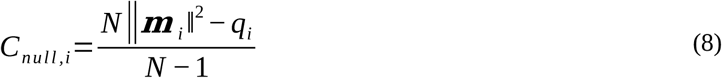

For the observed covariance at pair separation m, the sequence-excess covariance and its normalized form were

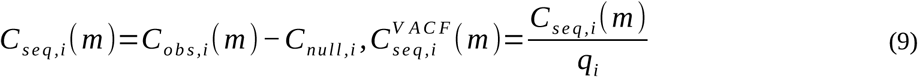

Observed covariance was required to equal the sum of the order-null and sequence-excess components to machine precision. The order-null term quantifies covariance attributable to static trajectory polarity; the sequence-excess term isolates serial organization beyond those preserved static properties.

### Exact displacement-memory decomposition

For exact finite-track closure, pair covariances were defined inside displacement windows of length n. For each separation m, the window-conditioned covariance was the average dot product over all increment pairs at that separation contained in all valid n-step windows. Then

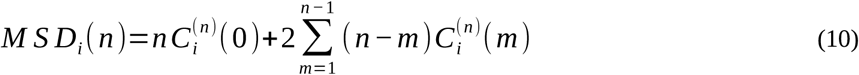

The diagonal term is activity and the off-diagonal terms are displacement memory:

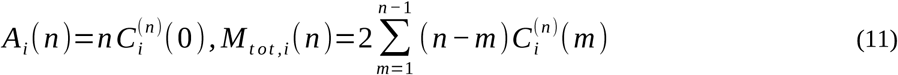

Replacing each distinct-pair covariance by the trajectory-specific permutation expectation yielded the order- null memory term. Sequence-excess memory was therefore

[math12]

Factorial interactions in direct displacement, activity, order-null memory, and sequence-excess memory were normalized by the repeat-specific shCTRL direct MSD at the same displacement lag. All numerical outputs were rejected unless direct=activity+order-null+sequence-excess closed to floating-point tolerance.

### Pair-separation bridge and kernel-opportunity correction

For a source set of pair separations, only members shorter than the n-step displacement window were eligible. This n-specific subset is denoted in Equations 13 and 14. The sequence-excess contribution of that source set to n-step displacement memory was

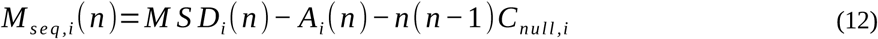

The source band was sequentially frozen as m=2,…,12 (20–120 min) after the sequence-excess VACF analysis and before evaluating the bridge; the target displacement window was n=12,…,20 (120–200 min). Because the triangular kernel offers unequal numbers of pair opportunities to different separations, the exact opportunity and density were

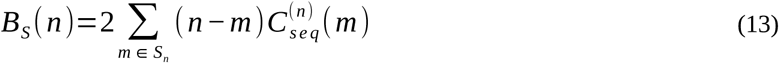

Preferential enrichment was tested by subtracting the opportunity-normalized density outside the source set from that within the source set. A positive within-source density establishes positive contribution per available pair opportunity; a positive within-minus-outside contrast would be required to claim preferential concentration in the source set.

### Exact speed–direction decomposition

Write each increment as xᵢ = sᵢeᵢ, where sᵢ = ‖xᵢ‖ is its speed magnitude and eᵢ is its unit direction. Pairwise directional alignment is aᵢⱼ = eᵢ·eⱼ. At a given lag, the exact expectation identity is

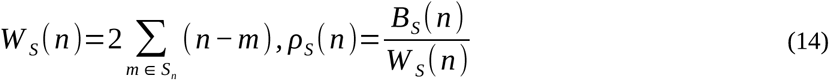

The three terms quantify mean-speed directional persistence, speed-memory modulation, and speed– direction coupling. Each term was computed for every trajectory and lag, propagated through the FOV– repeat hierarchy, and expressed as a factorial interaction normalized by repeat-specific shCTRL mean squared increment. Algebraic closure was checked at the FOV and interaction levels.

### Collective-flow decomposition and leave-one-cell-out estimator

For each focal trajectory and time point, the primary shared-flow estimator was the arithmetic mean of all contemporaneous trajectories in the same FOV except the focal trajectory; it was defined whenever at least one other trajectory was present. The focal cell-relative increment was the focal increment minus this leave-one-cell-out field mean. The original inclusive FOV mean was retained as a sensitivity estimator. Inclusive and leave-one-cell-out decompositions were evaluated on identical lag-pair support so that their difference isolated focal-cell self-inclusion rather than occupancy-dependent missingness. In Equation 16, V denotes the leave-one-cell-out field mean and u the corresponding focal cell-relative increment.

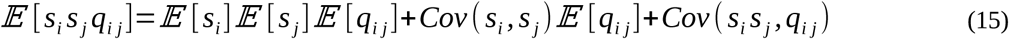

The terms are shared-flow covariance, cell-relative covariance, and two directional cross terms; the cross terms were summed for inference. Trajectory estimates were averaged within FOV, FOVs equally within repeat-condition strata, and R1–R3 equally. Exact closure was checked at FOV and interaction levels.

Polar order and distance-binned residual-velocity spatial correlations were retained as descriptive spatial summaries.

### Measured physical-context controls

FOV summaries included median occupancy, median nearest-neighbor distance, median image-edge distance, and median trajectory duration. Absolute coordinates passed a coordinate-spread audit before spatial analyses were enabled. Within each biological repeat, shCTRL FOVs anchored four-condition quartets. FOVs in the other conditions were assigned without replacement using minimum-cost Hungarian matching after log1p transformation and standardization of the selected context variables.

Separate scenarios matched occupancy, density, edge distance, duration, joint nonspatial variables, or all context variables. A direct edge analysis excluded increment pairs whose positions were <20 μm from the image boundary. A duration sensitivity required at least 48 observations. These analyses test robustness to the measured variables; they do not constitute randomized control of unmeasured FOV biology.

### Cross-system TrackMate validation and archive-cohort sensitivity

We independently evaluated the frozen serial-order framework using the public TrackMate XML resource of LaChance et al. (2021, 2022). XML spot coordinates were linked along TrackMate edges, nonconsecutive temporal segments were split, and the detected time interval was checked against the source design. The archive supplied 17 MDA-MB-231 XML movies, 18 HUVEC XML exports, 15 MDCK-bulk XMLs, and 16 MDCK-edge XMLs. The publisher-labelled MDCK-edge file 01_Tracks_BAD.xml was excluded before analysis, leaving 65 movies. Sampling intervals were 5 min for MDA-MB-231 and 10 min for HUVEC and MDCK. A trajectory entered the fixed cohort only if a contiguous segment remained observable through 120 min. XML/movie was the inferential unit; trajectories were weighted equally within movie and movies equally within system.

Primary endpoints were observed normalized increment covariance, the analytical within-trajectory order- null, sequence excess, sequence-excess area under the curve (AUC) over 20–120 min, the last lag with a simultaneously positive sequence-excess band, the frozen 60-min checkpoint, and the exact 60-min decomposition into activity, order-null memory, and serial-order memory. Within each system, 10,000 movie-level bootstrap replicates (seed 20260716) generated pointwise intervals and maximum-deviation simultaneous bands. Because species, substrate, acquisition, and native spatial units differ, cross-system contrasts were interpreted descriptively; native speeds and spatial magnitudes were not compared.

The source paper reports 13 HUVEC movies in its modelling workflow, whereas the public archive contains 18 valid XML exports. All 18 exports therefore defined the primary archive cohort, and all 8,568 possible 13- of-18 subsets were evaluated without selecting a preferred subset. MDCK bulk and edge were compared only for 14 identical filename-derived tissue keys, conservatively treated as putative matches because dish

metadata were unavailable. Unmatched files remained in unpaired system summaries. Four prespecified paired MDCK endpoints were compared using two-sided Wilcoxon signed-rank tests. Familywise error across those four tests was controlled with a Bonferroni threshold of α=0.0125, and exact P values are reported. Four-system omnibus tests were excluded from manuscript inference because MDCK bulk and edge are paired and major experimental covariates differ among systems.

### Single-cell haptotaxis dataset and locked closure analysis

Complete single-cell trajectories were obtained from the public data accompanying Fortunato et al. (2025; data DOI 10.34810/data2489). The analyzed cohort contained 131 MCF10A trajectories sampled at 10-min intervals on fibronectin gradients confined to 20, 40, 60, 80, or 250 µm. Deposited source labels were used to preserve the recoverable experimental-block hierarchy. The primary geometry contrast compared 250 µm with the arithmetic mean of 20–80 µm and used equal-block hierarchical bootstrap intervals.

For each track, sequence-dependent persistence was resolved along the fibronectin-gradient axis and the transverse axis. Axis contribution used the total activity denominator and therefore combines activity allocation with intrinsic order. Intrinsic axis order divided the axis-specific sequence excess by activity on the same axis, separating how often a cell moved laterally from how ordered those lateral movements were. Frozen windows were 20–60, 20–120, and 60–180 min. The finalized closure analysis directly bootstrapped the early-minus-late contrast and tested the 6-h-minus-3-h observation-horizon difference.

Reversal events were detected from sustained changes in gradient-axis direction. The finalized event analysis used the same complete-case cells for pre-transition, transition, and post-transition windows. Matched pseudo-events were sampled within the same tracks while preserving the event-time and available-duration distributions. The true reversal statistic was compared with the pseudo-event distribution to prevent a direction switch selected by construction from being interpreted as a reversal-specific biological peak.

### PFKL chemotaxis dataset and locked post hoc robustness analysis

Tracking CSVs were obtained from Hansen and Webb (2025) and Dryad dataset 10.5061/dryad.6m905qgfp. MDA-MB-231 cells were imaged every 10 min for 16 h in an EGF gradient directed along negative x. Fourteen tracking files contributed 1,691 reconstructed tracks. The primary comparison was PFKL-WT versus filament-incompetent PFKL-N702T (150 tracks per group). Secondary within-study comparisons were siPFKL versus siCTRL, shPFKL versus shScramble, PFKL-H199Y versus PFKL-WT, and PFK15 versus DMSO. The deposited CSVs did not retain biological-replicate identity; inference was therefore explicitly track-level.

Conventional endpoints included mean speed, cue-axis and transverse forward migration index, final cue displacement, path persistence, track duration, and missing-frame structure. Temporal endpoints included total order, cue-axis contribution, intrinsic cue-axis order, up-gradient step fraction, cue-sign persistence, switching rate, and mean and maximum up-gradient run length. The PFKL analysis used the same analytical order-null principle but did not apply discovery-dataset factorial inference.

After exploratory Stage 1 and Stage 2 audits, a post hoc Stage 3 analysis plan was fixed before final execution. The primary HC3 regression was outcome ∼ N702T + log(mean speed) + duration. Number of positions was excluded from the primary model because it correlated 0.999988 with duration and was used only in a separate sensitivity model. Optimal 1:1 propensity matching without replacement used log speed and duration, a caliper of 0.20 SD of the propensity-score logit, a minimum of 100 retained pairs, and a maximum post-match absolute standardized mean difference below 0.10. Overlap weighting, pooled speed-quartile analyses, common-duration support, split-track exclusion, cue-activity restriction, and speed- outlier exclusion were additional locked sensitivities.

For intrinsic cue-axis order, total serial order, and cue-sign persistence, track-level non-inferiority was evaluated against a −0.10-SD margin. Non-inferiority was concluded when the one-sided 95% lower confidence bound for the standardized N702T effect exceeded −0.10. Bootstrap tail proportions were retained only as diagnostics and were not interpreted as formal non-inferiority P values.

### Evidence freeze before generative modelling

The empirical evidence ledger was frozen after the reconstruction, factorial, order-null, memory-spectrum, kernel, speed–direction, collective-mode, and context analyses. Frozen claims, unsupported claims, lag windows, source tables, and figure-panel mappings were written before the GPU models were evaluated. The generative models therefore could not redefine successful observables or lag intervals after seeing their predictive performance. The external-validation specification, including the 20–120-min AUC, 60-min checkpoint, 120-min fixed cohort, movie-level hierarchy, and archive-discrepancy sensitivity, was written before external results were inspected.

### Stationary linear active-memory model

The first generative model was fitted only to local one- and two-step transitions. For condition-repeat group g, shared FOV flow followed a vector AR(1) process

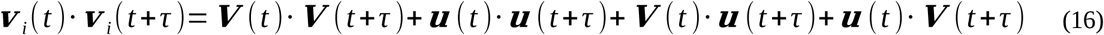

Cell-relative increments followed a stable vector AR(2) process driven by the preceding flow level and flow change:

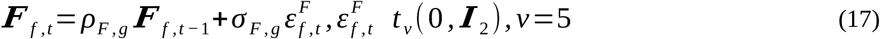

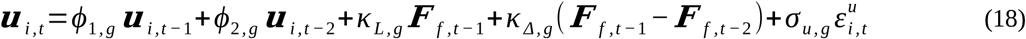

AR(2) stability was enforced through partial-autocorrelation variables π₁ and π₂:

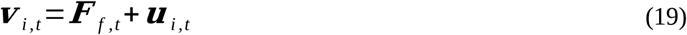

Parameters were bounded by tanh transforms and positive scales by softplus transforms. With residual e in either coordinate, the fixed-degree-of-freedom Student-t negative log-likelihood, up to constants common to all fits, was

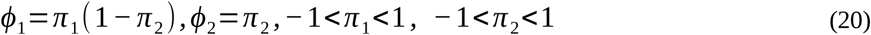

Sixteen FOV-bootstrap models were optimized simultaneously. The objective was residual-transition loss plus 0.5 times the flow-transition loss and a small quadratic raw-parameter penalty. Each model generated eight complete experiments after a 60-frame burn-in, preserving FOV counts, trajectory lengths, start times, and censoring. The no-interaction counterfactual replaced the combined-condition unconstrained parameter vector a₁₁ by

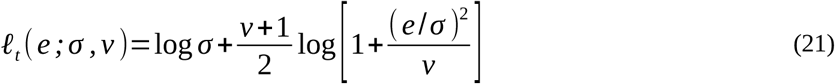

Additional ablations used one residual memory mode, removed flow–cell coupling, or removed shared flow. Neither MSD nor any frozen long-lag correlation or memory curve entered the fitting loss.

### Nonlinear angular hidden-state model with FOV random effects

The second model targeted angular organization that remained unexplained by the linear model. It conditioned on measured FOV-flow time series, measured cell-relative step magnitudes, trajectory start times, and trajectory lengths, and simulated hidden states and headings. Let θ denote cell-relative heading, φ the FOV-flow heading, α their wrapped angular difference, and Δφ the wrapped change in FOV-flow heading. Flow strength was transformed by the hyperbolic tangent of flow magnitude divided by the FOV median non-zero flow magnitude.

The hidden state z∈{P,R} represented a concentrated persistent state and a broader reorientation state. Conditional turning followed a von Mises density (Rabiner, 1989; Mardia and Jupp, 2000):

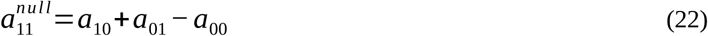

The state-specific mean turn incorporated nonlinear alignment and signed FOV turning:

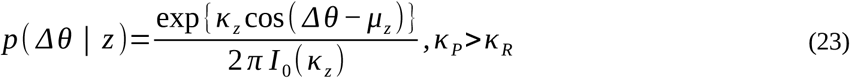

Switching probabilities were

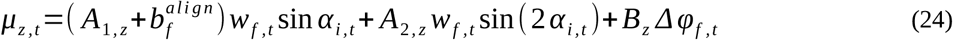

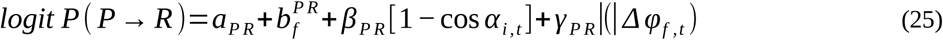

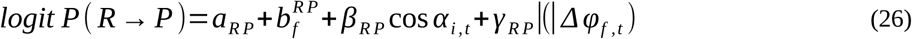

Gaussian-penalized FOV random effects represented between-FOV heterogeneity in both switching probabilities and alignment. State-label switching was prevented by parameterizing the persistent-state concentration as the reorientation-state concentration plus a positive gap. Likelihood was computed by the log-domain forward algorithm (Rabiner, 1989). The fit used inverse trajectory-count weighting within FOV so that densely tracked FOVs did not dominate. Two deterministic held-out FOVs per repeat–condition stratum (24 total) were evaluated with all FOV random effects set to zero. Ablations removed hidden-state switching, the second angular harmonic, FOV random effects, or alignment-dependent switching. Von Mises simulation used a vectorized Best–Fisher rejection sampler (Best and Fisher, 1979).

### Frozen predictive validation

For each scenario, 16 FOV-bootstrap models × 8 simulations generated 128 synthetic experiments. Each synthetic experiment was processed through the same exact VACF, order-null, speed–direction, collective-mode, sequence-memory, and bridge calculations as the observed data. Frozen windows were 30–150 min for VACF, 20–120 min for sequence-excess VACF, 40–100 min for sequence-excess covariance, 50–150 min for the directional term, 80–140 min for shared flow, 70–150 min for cell-relative motion, 10–160 min for the negative cross term, and 120–200 min for sequence memory and the bridge.

A predictive target passed only when at least 80% of its lag points were covered by the 95% predictive interval, the observed frozen-window mean lay within the corresponding predictive interval, and the probability of the correct mean sign was at least 0.90. Sign-hierarchy frequency was reported secondarily and was not used to override quantitative failure.

### Software, computation, and reproducibility

Empirical analyses were run as versioned Python/SLURM workflows on the Hummel-2 cluster. CPU jobs requested Hummel-2-compatible multiples of eight cores; GPU model fitting used one NVIDIA H100 80-GB GPU and eight CPUs. Exact cumulative-sum identities replaced memory-intensive displacement-window tensors. Each workflow writes input hashes, frozen configurations, random seeds, source-data tables, numerical checks, software versions, and a compressed result archive. A complete peer-review package accompanies this appeal and contains the discovery workflow, haptotaxis closure, PFKL Stage 3 robustness analysis, command-line track workflow, deterministic tests, source tables, software documentation, frozen configurations, numerical audits, and figure-rendering code. OpenAI ChatGPT was used for language editing, document organization, code review, and figure-layout refinement; it was not used to generate source data or experimental measurements. All numerical results, code, citations, and interpretations were checked by the author, who takes full responsibility for the work.

## DATA AVAILABILITY

The MYO10-collagen discovery trajectories were generated for Peuhu et al. (2022), compiled in CellTracksColab (Gómez-de-Mariscal et al., 2024), and deposited on Zenodo (doi:10.5281/zenodo.11282716; Jacquemet et al., 2024). External MDA-MB-231, HUVEC, and MDCK TrackMate files are available from Zenodo (doi:10.5281/zenodo.4959169; LaChance et al., 2021, 2022). Single-cell haptotaxis data are available from the CORA Research Data Repository (doi:10.34810/data2489; Fortunato et al., 2025). PFKL chemotaxis data are available from Dryad (doi:10.5061/dryad.6m905qgfp; Hansen and Webb, 2025). The accompanying peer-review package contains corrected source tables, figure-rendering code, software documentation, frozen configurations, input manifests, numerical audits, and the haptotaxis and PFKL result archives. A DOI-minted public software and results repository will be deposited before publication.

## AUTHOR CONTRIBUTIONS

Subhajit Dutta: Conceptualization; Methodology; Software; Validation; Formal analysis; Investigation; Data curation; Visualization; Writing – original draft; Writing – review and editing; Project administration.

## COMPETING INTERESTS

The author declares no competing interests.

## ACKNOWLEDGMENTS

The author thanks E. Peuhu, G. Jacquemet, C.L.G.J. Scheele, the CellTracksColab contributors, J. LaChance, K. Suh, D.J. Cohen, I.C. Fortunato, D.B. Brückner, X. Trepat, H.L. Hansen, B.A. Webb, and the other original investigators and data curators whose public datasets made this reanalysis possible. These acknowledgments do not imply endorsement of the present analysis or conclusions.

The Hummel-2 high-performance computing cluster at Universität Hamburg was used for this work. The cluster was funded by the Deutsche Forschungsgemeinschaft (DFG, German Research Foundation) under project number 478636198.

## Notes

### Competing Interest Statement

The authors have declared no competing interest.

### Summary of Updates

This version has been substantially revised to improve biological framing, accessibility, methodological clarity, and practical usability. The title has been changed to “Temporal order separates generic persistence from cue-directed commitment in cell migration.” The Introduction and Results have been reorganized around the biological question of whether temporal ordering can distinguish general persistence from productive alignment to an environmental cue. A new introductory schematic and a plain-language glossary have been added to explain the main concepts, including the order-null reference, sequence-dependent persistence, accumulated displacement memory, factorial interaction, and cue-aligned commitment. The statistical and modelling sections have been streamlined and moved after the principal biological findings. New analyses have been added for single-cell haptotaxis in confined fibronectin gradients and for PFKL-dependent chemotaxis toward EGF. These analyses show that physical confinement redistributes temporal order between spatial axes, while PFKL-N702T impairs cue-aligned migration without a resolved loss of mean speed or generic serial order. Cross-system evaluation across endothelial, epithelial, and cancer-cell datasets has also been expanded. All main and supplementary figures have been redesigned, renumbered, and updated. The Appendix now includes expanded mathematical derivations, numerical audits, robustness analyses, matching and weighting diagnostics, model-validation results, and source-controlled supplementary figures. A documented user-facing workflow has been added for calculating conventional and temporal-order migration measures from standard tracking tables. Limitations concerning biological replication, public metadata, cross-system comparisons, and causal interpretation have been clarified throughout.

## REFERENCES

1. Best, D.J., and N.I. Fisher. 1979. Efficient simulation of the von Mises distribution. Appl. Stat. 28:152–157. doi:10.2307/2346732.

2. Bohil, A.B., B.W. Robertson, and R.E. Cheney. 2006. Myosin-X is a molecular motor that functions in filopodia formation. Proc. Natl. Acad. Sci. USA. 103:12411–12416. doi:10.1073/pnas.0602443103.

3. Efron, B., and R.J. Tibshirani. 1993. An Introduction to the Bootstrap. Chapman & Hall, New York.

4. Fortunato, I.C., D.B. Brückner, S. Grosser, et al. 2025. Single-cell migration along and against confined haptotactic gradients. Nat. Phys. 21:1638–1647. doi:10.1038/s41567-025-03015-3.

5. Gorelik, R., and A. Gautreau. 2014. Quantitative and unbiased analysis of directional persistence in cell migration. Nat. Protoc. 9:1931–1943. doi:10.1038/nprot.2014.131.

6. Gómez-de-Mariscal, E., H. Grobe, J.W. Pylvänäinen, et al. 2024. CellTracksColab is a platform that enables compilation, analysis, and exploration of cell tracking data. PLoS Biol. 22:e3002740. doi:10.1371/journal.pbio.3002740.

7. Hansen, H.L., and B.A. Webb. 2025. Functional requirements of the liver isoform of phosphofructokinase-1 in breast cancer cell migration. J. Cell Sci. 138:jcs264251. doi:10.1242/jcs.264251.

8. Jacquemet, G., E. Gómez-de-Mariscal, H. Grobe, J.W. Pylvänäinen, L. Xénard, and R. Henriques. 2024. CellTracksColab - breast cancer cell dataset. Zenodo. doi:10.5281/zenodo.11282716.

9. LaChance, J., K. Suh, and D.J. Cohen. 2021. Deep attention networks for automated collective behavior discovery in epithelia. Zenodo. doi:10.5281/zenodo.4959169.

10. LaChance, J., K. Suh, J. Clausen, and D.J. Cohen. 2022. Learning the rules of collective cell migration using deep attention networks. PLoS Comput. Biol. 18:e1009293. doi:10.1371/journal.pcbi.1009293.

11. Licup, A.J., S. Münster, A. Sharma, et al. 2015. Stress controls the mechanics of collagen networks. Proc. Natl. Acad. Sci. USA. 112:9573–9578. doi:10.1073/pnas.1504258112.

12. Maiuri, P., J.-F. Rupprecht, S. Wieser, et al. 2015. Actin flows mediate a universal coupling between cell speed and cell persistence. Cell. 161:374–386. doi:10.1016/j.cell.2015.01.056.

13. Mardia, K.V., and P.E. Jupp. 2000. Directional Statistics. Wiley, Chichester, UK.

14. Miihkinen, M., M.L.B. Grönloh, A. Popović, et al. 2021. Myosin-X and talin modulate integrin activity at filopodia tips. Cell Rep. 36:109716. doi:10.1016/j.celrep.2021.109716.

15. Paszek, M.J., N. Zahir, K.R. Johnson, et al. 2005. Tensional homeostasis and the malignant phenotype. Cancer Cell. 8:241–254. doi:10.1016/j.ccr.2005.08.010.

16. Peuhu, E., G. Jacquemet, C.L.G.J. Scheele, et al. 2022. MYO10-filopodia support basement membranes at pre-invasive tumor boundaries. Dev. Cell. 57:2350–2364.e7. doi:10.1016/j.devcel.2022.09.016.

17. Popović, A., M. Miihkinen, S. Ghimire, et al. 2023. Myosin-X recruits lamellipodin to filopodia tips. J. Cell Sci. 136:jcs260574. doi:10.1242/jcs.260574.

18. Provenzano, P.P., D.R. Inman, K.W. Eliceiri, et al. 2008. Collagen density promotes mammary tumor initiation and progression. BMC Med. 6:11. doi:10.1186/1741-7015-6-11.

19. Rabiner, L.R. 1989. A tutorial on hidden Markov models and selected applications in speech recognition. Proc. IEEE. 77:257–286. doi:10.1109/5.18626.

20. SenGupta, S., C.A. Parent, and J.E. Bear. 2021. The principles of directed cell migration. Nat. Rev. Mol. Cell Biol. 22:529–547. doi:10.1038/s41580-021-00366-6.

21. Skoge, M., H. Yue, M. Erickstad, et al. 2014. Cellular memory in eukaryotic chemotaxis. Proc. Natl. Acad. Sci. USA. 111:14448–14453. doi:10.1073/pnas.1412197111.

22. Westfall, P.H., and S.S. Young. 1993. Resampling-Based Multiple Testing: Examples and Methods for p-Value Adjustment. Wiley, New York.

23. Wolf, K., M. Te Lindert, M. Krause, et al. 2013. Physical limits of cell migration: control by ECM space and nuclear deformation and tuning by proteolysis and traction force. J. Cell Biol. 201:1069–1084. doi:10.1083/jcb.201210152.

