## Appendix_1 for "Temporal order separates generic persistence from cue-directed commitment in cell migration"

Temporal order separates generic persistence from
cue-directed commitment in cell migration

**Subhajit Dutta**

Institute of Biochemistry and Molecular Cell Biology, Center for Experimental Medicine, University Medical Center Hamburg-Eppendorf, Hamburg, Germany

#### CONTENTS

Appendix 1—note 1. Notation and hierarchy

Appendix 1—note 2. Step 1–17 computational workflow

Appendix 1—note 3. Exact temporal-order null

Appendix 1—note 4. Finite-trajectory displacement-memory derivation

Appendix 1—note 5. Pair-separation kernel and enrichment test

Appendix 1—note 6. Speed–direction and collective-flow identities

Appendix 1—note 7. Hierarchical bootstrap and robustness gates

Appendix 1—note 8. Physical-context matching

Appendix 1—note 9. Linear active-memory model

Appendix 1—note 10. Nonlinear angular hidden-state model

Appendix 1—note 11. Predictive-validation design

Appendix 1—note 12. Interpretation boundaries and reproducibility

Appendix 1—note 13. Leave-one-cell-out collective decomposition

Appendix 1—note 14. Independent external validation

Appendix 1—note 15. Single-cell haptotaxis analysis and closure

Appendix 1—note 16. PFKL chemotaxis analysis and inferential lock

Appendix 1—note 17. User-facing workflow and interpretation boundaries

Appendix 1 tables. Extended numerical summaries

Appendix 1 figures. Extended diagnostic figures

#### APPENDIX 1—NOTE 1. NOTATION AND HIERARCHY

For accessibility, the main text uses one canonical name for each concept: order-null for the exact shuffled-order expectation, sequence excess or sequence-dependent persistence for the observed-minus-null contrast, generic serial order for cue-independent continuation, axis contribution for spatial allocation, and intrinsic axis order for ordering normalized within one axis. Main-text Table 1 provides the full plain-language glossary. In the equations below, indices i, f, r, g, t, m, and n denote trajectory, FOV, biological repeat, repeat-condition group, time origin, pair separation, and displacement-window length, respectively. The sampling interval is $\Delta t$=10 min. Condition indices 00, 01, 10, and 11 denote shCTRL, shCTRL+collagen, shMYO10, and shMYO10+collagen. All primary empirical observables follow the aggregation order trajectory → FOV → biological repeat; the repeat mean gives equal weight to R1, R2, and R3.

$$y_{r,c}^{rep}=\frac{1}{card(F_{r,c})}\sum_{f\in F_{r,c}}\frac{1}{card(I_f)}\sum_{i\in I_f}y_i,\; y_c^{cond}=\frac{1}{3}\sum_{r=1}^{3}y_{r,c}^{rep} \tag{S1}$$

The formula is schematic: the trajectory-level quantity may itself be an average over valid origins. Equal FOV weighting is retained even when FOVs contain different numbers of tracked identities.

#### APPENDIX 1—NOTE 2. STEP 1–17 COMPUTATIONAL WORKFLOW

The analysis was deliberately staged so that identity correction and empirical exact decompositions preceded all generative modelling. Appendix 1—table 1 gives the complete workflow. Each step wrote a dedicated result directory, source-data tables, run metadata, input hashes, and a compressed archive. Step 14 froze the evidence hierarchy before Steps 15–17 extended the physical decomposition and modelling.

#### APPENDIX 1—NOTE 3. EXACT TEMPORAL-ORDER NULL

Let a trajectory comprise N increments $x_1,\ldots,x\_N$. Define the mean increment vector $m = N^{-1}\Sigma_i x_i$ and the mean squared increment magnitude $q = N^{-1}\Sigma_i \|x_i\|^2$, and let π be a uniformly random permutation. For two distinct positions, all ordered pairs of distinct original increments are equally likely. Therefore

$$\mathbb{E}_\pi[\boldsymbol{x}_{\pi(a)}\cdot\boldsymbol{x}_{\pi(b)}]=\frac{1}{N(N-1)}\sum_{i\neq j}\boldsymbol{x}_i\cdot\boldsymbol{x}_j \tag{S2}$$

The squared norm of the increment sum equals the sum of the squared increment norms plus all distinct cross terms. Substitution of m and q gives

$$\sum_{i\neq j}\boldsymbol{x}_i\cdot\boldsymbol{x}_j=N^2\|\boldsymbol{m}\|^2-Nq \tag{S3}$$

$$C_{null}=\frac{N\|\boldsymbol{m}\|^2-q}{N-1} \tag{S4}$$

The null is analytical; no Monte Carlo permutations are required. It preserves the entire set of increments and hence step-length distribution, orientation distribution, static mean vector, total displacement, and trajectory length. It removes only their temporal arrangement. For observed pair covariance at separation m,

$$C_{obs}(m)=\frac{1}{N-m}\sum_{t=1}^{N-m}\boldsymbol{x}_t\cdot\boldsymbol{x}_{t+m},C_{seq}(m)=C_{obs}(m)-C_{null} \tag{S5}$$

$$VACF_{obs}(m)=\frac{C_{obs}(m)}{q},VACF_{null}=\frac{C_{null}}{q},VACF_{seq}(m)=\frac{C_{seq}(m)}{q} \tag{S6}$$

Because the order-null value is trajectory specific, normalization and averaging were performed at trajectory level before hierarchical aggregation. This prevents trajectories with large mean squared increments from dominating the normalized sequence-excess estimate.

#### APPENDIX 1—NOTE 4. FINITE-TRAJECTORY DISPLACEMENT-MEMORY DERIVATION

For a valid n-step window beginning at time origin t, let the cumulative increment over that window be denoted by ΔR(n,t). Expanding its squared norm gives

$$\|\Delta\boldsymbol{R}_t(n)\|^2=\sum_{j=0}^{n-1}\|\boldsymbol{x}_{t+j}\|^2+2\sum_{m=1}^{n-1}\sum_{j=0}^{n-m-1}\boldsymbol{x}_{t+j}\cdot\boldsymbol{x}_{t+j+m} \tag{S7}$$

Averaging over all valid origins yields an exact finite-track identity. Define

$$C^{(n)}(0)=\frac{1}{n(N-n+1)}\sum_t\sum_{j=0}^{n-1}\|\boldsymbol{x}_{t+j}\|^2 \tag{S8}$$

$$C^{(n)}(m)=\frac{1}{(n-m)(N-n+1)}\sum_{t}\sum_{j=0}^{n-m-1}\boldsymbol{x}_{t+j}\cdot\boldsymbol{x}_{t+j+m} \tag{S9}$$

$$MSD(n)=nC^{(n)}(0)+2\sum_{m=1}^{n-1}(n-m)C^{(n)}(m) \tag{S10}$$

This definition avoids an implicit stationarity approximation because the covariance at each pair separation is conditioned on the same displacement windows used for the n-step MSD. The components are

$$A(n)=nC^{(n)}(0),M_{tot}(n)=MSD(n)-A(n) \tag{S11}$$

$$M_{null}(n)=n(n-1)C_{null},M_{seq}(n)=M_{tot}(n)-M_{null}(n) \tag{S12}$$

$$MSD(n)=A(n)+M_{null}(n)+M_{seq}(n) \tag{S13}$$

The implementation also calculated pair-separation contributions directly using cumulative sums of increment dot products at each separation. Machine-precision closure was required at the trajectory, FOV, repeat, and factorial-interaction levels.

#### APPENDIX 1—NOTE 5. PAIR-SEPARATION KERNEL AND ENRICHMENT TEST

For a source set of pair separations, only members shorter than the n-step displacement window were eligible. The resulting n-specific subset contributes to displacement memory as

$$B_S(n)=2\sum_{m\in S_n}(n-m)C_{seq}^{(n)}(m) \tag{S14}$$

The coefficient 2(n−m) is the triangular displacement kernel. It counts ordered symmetric cross terms for pair separation m inside a window of n increments. The corresponding opportunity is

$$W_S(n)=2\sum_{m\in S_n}(n-m) \tag{S15}$$

$$\rho_S(n)=\frac{B_S(n)}{W_S(n)} \tag{S16}$$

For the source separations corresponding to 20–120 min and target windows corresponding to 120–200 min, the source opportunity fraction ranged from 0.833 to 0.753, with arithmetic mean approximately 0.813. The raw source share of the positive sequence contribution was approximately 0.872. Their difference is descriptive only. Inference was performed on the density difference

$$E(n)=\rho_S(n)-\rho_{S^c}(n) \tag{S17}$$

The opportunity-normalized source density was repeat consistent, whereas the source-minus-outside contrast was not. Accordingly, the source set contributes positively per opportunity but is not preferentially enriched relative to valid outside separations.

#### APPENDIX 1—NOTE 6. SPEED–DIRECTION AND COLLECTIVE-FLOW IDENTITIES

Write each increment as $x_i = s_i e_i$, where $s_i = \|x_i\|$ is its speed magnitude and $e_i$ is its unit direction. Define pairwise directional alignment as $a_{ij} = e_i \cdot e_j$:

$$\mathbb{E}[s_i s_j q_{ij}]=\mathbb{E}[s_i s_j]\mathbb{E}[q_{ij}]+Cov(s_i s_j,q_{ij}) \tag{S18}$$

$$\mathbb{E}[s_i s_j]=\mathbb{E}[s_i]\mathbb{E}[s_j]+Cov(s_i,s_j) \tag{S19}$$

$$\mathbb{E}[s_i s_j q_{ij}]=\mathbb{E}[s_i]\mathbb{E}[s_j]\mathbb{E}[q_{ij}]+Cov(s_i,s_j)\mathbb{E}[q_{ij}]+Cov(s_i s_j,q_{ij}) \tag{S20}$$

The first term changes when mean speeds or directional persistence change; it was labeled mean-speed directional persistence. The second is speed-memory modulation, and the third is speed–direction coupling. The decomposition is algebraic and requires no probabilistic independence assumption.

For the primary collective decomposition, the field estimator for each focal cell was the arithmetic mean of all contemporaneous cells in the same FOV except that focal cell. The corresponding cell-relative increment was the focal increment minus this leave-one-cell-out field mean. Then

$$\boldsymbol{v}_i(t)\cdot\boldsymbol{v}_i(t+\tau)=C_F^{-i}+C_u^{-i}+C_{F\to u}^{-i}+C_{u\to F}^{-i} \quad \text{(S21)}$$

$$C_F^{-i}=\boldsymbol{V}_{f,-i}(t)\cdot\boldsymbol{V}_{f,-i}(t+\tau),C_u^{-i}=\boldsymbol{u}_{i,-i}(t)\cdot\boldsymbol{u}_{i,-i}(t+\tau) \quad \text{(S22)}$$

$$C_{F\to u}^{-i}=\boldsymbol{V}_{f,-i}(t)\cdot\boldsymbol{u}_{i,-i}(t+\tau),C_{u\to F}^{-i}=\boldsymbol{u}_{i,-i}(t)\cdot\boldsymbol{V}_{f,-i}(t+\tau) \quad \text{(S23)}$$

Unlike residuals defined around the inclusive mean, leave-one-cell-out residuals do not sum to zero over focal cells at each time. This does not affect the exact per-focal-cell identity. The two lagged cross terms can be nonzero and were summed for inference. The inclusive decomposition was retained on identical support as a sensitivity analysis.

#### APPENDIX 1—NOTE 7. HIERARCHICAL BOOTSTRAP AND ROBUSTNESS GATES

Let the observed repeat-equal-weighted interaction curve be denoted by J_obs and the b-th hierarchical bootstrap curve by J_b. Pointwise intervals were empirical 2.5th and 97.5th percentiles. For simultaneous inference, define the bootstrap standard error s(τ) and

$$T_b=max_{\tau\in G}\frac{\|[J_b(\tau)-J_{obs}(\tau)]\|}{s(\tau)} \quad \text{(S24)}$$

$$Band(\tau)=J_{obs}(\tau)\pm q_{0.95}(T)s(\tau) \quad \text{(S25)}$$

The robust-window gate additionally required a common repeat sign and a leave-one-repeat-out sign check: for each omitted repeat, the mean of the two retained repeats had to have the same sign as the omitted repeat. This conservative conjunction was chosen because n=3 at the highest level. Bootstrap intervals quantify stability under the observed hierarchy; they do not create additional biological replicates.

#### APPENDIX 1—NOTE 8. PHYSICAL-CONTEXT MATCHING

For each FOV, the context vector contained selected combinations of log1p occupancy, nearest-neighbor distance, image-edge distance, and trajectory duration, standardized within the matching analysis. For an anchor control FOV and a candidate FOV, the matching cost was squared Euclidean distance

$$d(a,b)=\|\boldsymbol{z}(\boldsymbol{h}_a)-\boldsymbol{z}(\boldsymbol{h}_b)\|_2^2 \quad \text{(S26)}$$

Within each biological repeat, the Hungarian algorithm selected one-to-one assignments from each noncontrol condition to the shCTRL anchors. Matching was repeated for individual context dimensions and joint sets. Balance tables report before/after standardized differences. The primary inferential quantity remained the repeat-specific one-hour VACF interaction after restriction to matched FOV quartets.

#### APPENDIX 1—NOTE 9. LINEAR ACTIVE-MEMORY MODEL

The full model was condition- and repeat-specific. Shared flow and residual increments were two-dimensional with scalar autoregressive coefficients shared across coordinates. Innovations were independent standardized Student-t variables with ν=5 and condition-repeat-specific scales.

$$\boldsymbol{F}_t=\rho_F\boldsymbol{F}_{t-1}+\sigma_F\varepsilon_t^F \quad \text{(S27)}$$

$$\boldsymbol{u}_t=\phi_1\boldsymbol{u}_{t-1}+\phi_2\boldsymbol{u}_{t-2}+\kappa_L\boldsymbol{F}_{t-1}+\kappa_\Delta(\boldsymbol{F}_{t-1}-\boldsymbol{F}_{t-2})+\sigma_u\varepsilon_t^u \quad \text{(S28)}$$

$$\boldsymbol{v}_t=\boldsymbol{F}_t+\boldsymbol{u}_t \quad \text{(S29)}$$

Bounded transformations used 0.995 times a hyperbolic tangent for shared-flow persistence, 0.98 times a hyperbolic tangent for each partial-autocorrelation parameter, 1.5 times a hyperbolic tangent for each flow-coupling coefficient, and softplus plus $10^{-4}$ for positive scales. The partial-autocorrelation map places the AR(2) polynomial in the stationary region.

For each bootstrap ensemble, the multiplicity assigned to an observation's FOV weighted its likelihood contribution. The optimized objective was

$$L = mean_e\left[L_u^{(e)} + 0.5\, L_F^{(e)}\right] + 10^{-4}\, mean\left(a^2\right) \tag{S30}$$

Each L is a FOV-bootstrap-weighted mean of coordinate-summed Student-t NLL. Optimization used AdamW, cosine learning-rate decay, gradient clipping at 10, 8,000 steps for the full model, and 6,000 for ablations. OLS estimates initialized the unconstrained parameters. The full simulation used 60 burn-in frames, then retained the empirical FOV duration and trajectory censoring masks. Residual increments were re-centered at each time to reproduce the exact Step 15 decomposition.

The prespecified scenarios were the full model, a no-interaction counterfactual, a one-mode residual process, a no-coupling model, and a no-flow AR(2) process fitted directly to total increments. The no-interaction parameter for the combined condition was constrained to the additive expectation on the unconstrained scale; the one-mode scenario set the second residual coefficient to zero; and the no-coupling scenario set both flow-coupling coefficients to zero. Sixteen FOV-bootstrap ensembles and eight simulations per ensemble produced 128 experiments per scenario. Appendix 1—figure 8 presents endpoint-level posterior-predictive diagnostics for the related fractional active-memory model zoo; the frozen long-lag pass/fail results for the principal linear model remain in Appendix 1—table 6.

#### APPENDIX 1—NOTE 10. NONLINEAR ANGULAR HIDDEN-STATE MODEL

Residual headings were computed from the two Cartesian components of cell-relative increments, and FOV-flow headings from the two components of the field vector. Reliable angular emissions required adjacent residual step magnitudes of at least 0.10 µm. All 48,134 primary trajectories were retained in conditional simulation, while trajectories with at least three reliable emissions entered fitting.

$$\delta\theta_t = wrap\left(\theta_t - \theta_{t-1}\right), \alpha_t = wrap\left(\varphi_{t-1} - \theta_{t-1}\right), \delta\varphi_t = wrap\left(\varphi_{t-1} - \varphi_{t-2}\right) \tag{S31}$$

$$w_t = \tanh\left(\frac{\left\|\boldsymbol{F}_{t-1}\right\|}{s_f}\right), s_f = median_{t:\|\boldsymbol{F}_t\| ¿ 0}\left\|\boldsymbol{F}_t\right\| \tag{S32}$$

For state z∈{P,R},

$$p\left(\delta\theta_t \mid z_t = z\right) = \frac{\exp\left\{\kappa_z \cos\left[\delta\theta_t - \mu_{z,t}\right]\right\}}{2\pi I_0\left(\kappa_z\right)} \tag{S33}$$

$$\mu_{z,t} = \left(A_{1,z} + b_f^{align}\right) w_t \sin\alpha_t + A_{2,z}\, w_t \sin\left(2\alpha_t\right) + B_z\, \delta\varphi_t \tag{S34}$$

$$logit\ p_{PR,t} = a_{PR} + b_f^{PR} + \beta_{PR}\left(1 - \cos\alpha_t\right) + \gamma_{PR}|(|\delta\varphi_t) \tag{S35}$$

$$logit\ p_{RP,t} = a_{RP} + b_f^{RP} + \beta_{RP}\cos\alpha_t + \gamma_{RP}|(|\delta\varphi_t) \tag{S36}$$

State concentrations were constrained so that the persistent state exceeded the reorientation state by a positive gap, fixing the state labels. Random effects were bounded transforms of unconstrained FOV parameters. The initial persistent probability was logistic in a group-specific parameter.

For the forward probability, emission probability, and state-transition probability, the likelihood recursion was

$$a_t(z) = e_t(z) \sum_{z' \in \{P,R\}} a_{t-1}\left(z'\right) p_{z' z,t}, p\left(y_{1:T}\right) = \sum_z a_T(z) \tag{S37}$$

Computation used log-sum-exp stabilization. Two FOVs per repeat-condition group were withheld by deterministic hash. Held-out likelihood set random effects to zero. In fitting, each trajectory received bootstrap FOV multiplicity divided by the number of angular trajectories in its FOV, providing equal FOV weight. Length-stratified minibatches reduced padding.

$$L_{HMM} = -wmean[\log p(y)] + 0.02 P_{FOV} + 0.002 P_{repeat} + 10^{-4} mean(a^2) \quad \text{(S38)}$$

Here, wmean denotes the prescribed inverse-trajectory-count-weighted mean within FOV. The FOV term was a Gaussian MAP penalty on raw FOV effects; the repeat term shrank repeat-specific group parameters toward the same-condition mean without forcing equality. The full model used 10,000 optimization steps and the ablations 7,500. Conditional simulations preserved observed speed, FOV-flow series, start times, lengths, and masks, generated hidden states and turns, and re-centered residual vectors to zero FOV mean. Ablations removed the second state, second harmonic, FOV random effects, or alignment-dependent switching; the no-interaction simulation was generated from the full fit on the unconstrained parameter scale. Appendix 1—figure 9 summarizes held-out likelihood, persistent-state occupancy, predictive coverage by target and scenario, and complete sign-hierarchy frequencies.

#### APPENDIX 1—NOTE 11. PREDICTIVE-VALIDATION DESIGN

For each metric and its frozen lag set, a simulated scenario passed only if

$$\frac{card\{\tau \in G_k : J_{obs}(\tau) \in PI_{0.95}(\tau)\}}{card(G_k)} \geq 0.80 \quad \text{(S39)}$$

$$mean_{\tau \in G_k} J_{obs}(\tau) \in PI_{0.95}\left[mean_{\tau \in G_k} J_{sim}(\tau)\right] \quad \text{(S40)}$$

$$P\left(sign[mean_{\tau \in G_k} J_{sim}] = sign_{expected}\right) \geq 0.90 \quad \text{(S41)}$$

The complete sign hierarchy required the expected sign in all three biological repeats for every frozen target within one synthetic experiment. It was reported as a secondary descriptor because a model can reproduce signs while failing magnitude and lag structure, as occurred for both generative models.

#### APPENDIX 1—NOTE 12. INTERPRETATION BOUNDARIES AND REPRODUCIBILITY

Supported statements are restricted to the trajectory-level physical observables. The analysis does not establish a direct molecular MYO10–collagen feedback, a unique intrinsic memory timescale, thermodynamic entropy production, or universal non-Markovian dynamics. Positive factorial interaction under suppressive main effects is buffering/antagonism. Negative flow–cell cross covariance is a kinematic cross term, not evidence of biochemical compensation.

The pair-separation source and target windows were selected sequentially within the same dataset and are therefore discovery-to-confirmation stages within one analysis, not independent replication. The highest inferential level is three biological repeats. All exact identities were unit tested and closed at machine precision. Step 14 froze the claim ledger before generative modelling. GPU runs were accepted only after status 0, complete output, and archive creation.

#### APPENDIX 1—NOTE 13. LEAVE-ONE-CELL-OUT COLLECTIVE DECOMPOSITION

For each focal cell, the primary field velocity was the arithmetic mean of all other contemporaneous cells in its FOV, and the cell-relative increment was the focal increment minus that leave-one-cell-out field velocity. Inclusive and leave-one-cell-out decompositions used identical lag-pair support. Therefore, their difference measures focal-cell self-inclusion rather than occupancy-dependent missingness. Trajectories were averaged within FOV, FOVs equally within biological repeat, and R1–R3 equally.

At 60 min, leave-one-cell-out total, shared-flow, cell-relative, and cross interactions were 0.115765, 0.019295, 0.102297, and −0.005827. Simultaneous intervals were [−0.004958, 0.236487], [0.000272, 0.038318], [−0.003085, 0.207679], and [−0.009992, −0.001663], respectively. LOO minus inclusive-common differences were exactly 0 for total, $+1.4\times10^{-5}$ for shared flow, $+3.91\times10^{-4}$ for cell-relative motion, and $-4.06\times10^{-4}$ for the cross term. All FOVs retained paired support; maximum closure error was $1.17\times10^{-15}$ µm². The detectable allocation change is quantitatively minimal. Appendix 1—table 4 reports the corresponding repeat-level estimates and simultaneous intervals.

#### APPENDIX 1—NOTE 14. INDEPENDENT EXTERNAL VALIDATION

The public external archive (LaChance et al., 2021; LaChance et al., 2022) contained 17 MDA-MB-231, 18 HUVEC, 15 MDCK-bulk, and 16 MDCK-edge XML exports. The publisher-labelled MDCK-edge file 01_Tracks_BAD.xml was excluded. Fixed cohorts required observation through 120 min. The movie/XML was the inferential unit, and exact decomposition closed with maximum residual zero. Primary endpoints were sequence-excess AUC over 20–120 min, the simultaneous positive horizon, sequence excess at 60 min, and exact 60-min serial-order displacement memory.

The source article reports 13 HUVEC movies in its modelling workflow, while the archive contains 18 valid exports. The primary estimate uses all 18 exports. Every one of the 8,568 possible 13-of-18 subsets was evaluated without selecting a preferred subset. MDCK bulk and edge comparisons used 14 identical filename-derived tissue keys and are described as putative matched identifiers. Four prespecified paired endpoints were evaluated using two-sided Wilcoxon signed-rank tests with a Bonferroni familywise threshold of $\alpha=0.0125$. Four-system omnibus tests were excluded from manuscript inference because paired MDCK observations violate independence and system identity is confounded with species, substrate, acquisition, and laboratory. The system curves, positive horizons, exact 60-min decompositions, exhaustive HUVEC subset analysis, and filename-matched MDCK comparison are shown in Appendix 1—figure 10.

#### APPENDIX 1—NOTE 15. SINGLE-CELL HAPTOTAXIS ANALYSIS AND CLOSURE

The haptotaxis extension used 131 complete MCF10A trajectories from fibronectin gradients of width 20, 40, 60, 80, and 250 µm, sampled at 10-min intervals (Fortunato et al., 2025; data DOI 10.34810/data2489). The recoverable source-block hierarchy was retained. The primary geometry contrast compared 250 µm with the arithmetic mean of 20–80 µm under equal-block hierarchical bootstrap.

Sequence-dependent persistence was resolved along the fibronectin-gradient and transverse axes. An axis-contribution score uses total activity in the denominator and therefore combines directional activity allocation with the intrinsic order of movements on that axis. The intrinsic axis-normalized score instead divides axis-specific sequence excess by activity on the same axis. This distinction was added during the haptotaxis audit and fixed before the final closure execution; it prevents the wide-geometry lateral effect from being attributed solely to cells taking more lateral steps.

At 20–60 min, the wide-minus-confined contrast was +0.1643 (95% hierarchical interval 0.1025–0.2271) laterally, −0.1182 (−0.2222 to −0.0021) along the gradient, and +0.0461 (−0.0542 to 0.1502) for total sequence-dependent order. The full-track intrinsic lateral-order contrast was +0.2176 (0.1274–0.3171), and the fully adjusted contrast was +0.2159 (0.1277–0.3092). The direct early-minus-late contrast was +0.1538 (0.0745–0.2386). The 6-h-minus-3-h observation-horizon contrast remained unresolved; no discrete six-hour onset is claimed. Primary, adjusted, alternative-window, common-support, and conventional-metric sensitivity analyses are assembled in Appendix 1—figure 11.

The reversal analysis used a common complete-case cohort for pre-transition, transition, and post-transition windows. Matched pseudo-event times were sampled within the same tracks while preserving event-time and available-duration distributions. This control distinguishes reversal-associated reorganization from the temporal segregation mechanically induced by selecting a sign change. Result language is restricted to a transient reorganization of temporal order, not a generic increase in one-direction persistence.

#### APPENDIX 1—NOTE 16. PFKL CHEMOTAXIS ANALYSIS AND INFERENTIAL LOCK

The metastatic chemotaxis extension used the tracking CSVs deposited by Hansen and Webb (2025; Dryad DOI 10.5061/dryad.6m905qgfp). MDA-MB-231 cells migrated toward a negative-x EGF gradient for 16 h with 10-min sampling. Fourteen files yielded 1,691 reconstructed tracks. The primary PFKL-WT versus PFKL-N702T comparison contained 150 tracks per group; secondary comparisons covered acute depletion, stable depletion, catalytic-dead PFKL-H199Y, and PFK15 treatment.

The source article reports three biological replicates per condition, but replicate identity is not recoverable from the compiled tracking CSVs. PFKL intervals are therefore track-level. The five perturbation series are interpreted as mechanistically distinct comparisons from one study rather than independent studies or biological-replicate confirmation.

A Stage 3 post hoc robustness plan was fixed before its final execution after exploratory Stage 1–2 auditing. The primary HC3 model was outcome ~ N702T + log(mean speed) + duration. Number of positions was excluded because it correlated 0.999988 with duration. Optimal propensity matching used log speed and duration, a 0.20-SD logit caliper, one-to-one matching without replacement, at least 100 pairs, and maximum post-match $|SMD|<0.10$. The final match retained 127 pairs with maximum $|SMD| =0.0089$. Overlap weighting achieved maximum $|SMD|=3.5\times10^{-11}$. Covariate balance, locked sensitivity filters, and the propensity-model specification are shown in Appendix 1—figure 12A,C,F.

N702T-minus-WT cue-FMI effects were −0.2783 unadjusted (track-bootstrap interval −0.3575 to −0.2000), −0.2797 in the primary HC3 model (−0.3580 to −0.2014), −0.2893 in matched pairs (−0.3710 to −0.2053), and −0.2797 after overlap weighting (−0.3573 to −0.2021). Mean up-gradient-run effects were −0.6663 steps unadjusted (−1.1275 to −0.2039), −0.5289 adjusted (−0.9774 to −0.0803), −0.6527 matched (−1.1581 to −0.1641), and −0.5285 weighted (−0.9910 to −0.0660). Mean speed was unresolved unadjusted and after matched comparison. The primary, matched, weighted, speed-stratified, and cross-series directional effects are summarized in main-text Figure 5 and Appendix 1—figure 12B,D.

Track-level non-inferiority used a locked −0.10-SD margin. One-sided 95% lower bounds were −0.0188 SD for intrinsic EGF-axis order, −0.0233 SD for total serial order, and −0.0314 SD for cue-sign persistence. All exceeded the margin. Bootstrap tail proportions were diagnostic only and are not reported as formal non-inferiority P values. Adjustment and matching support “not explained by measured speed differences,” not causal independence from speed. The non-inferiority estimates and one-sided lower bounds are shown in Appendix 1—figure 12E.

#### APPENDIX 1—NOTE 17. USER-FACING WORKFLOW AND INTERPRETATION BOUNDARIES

The portable workflow accepts delimited track tables with track identifier, time, x, and y, plus optional movie, condition, repeat, and cue-axis labels. It reports identifier reuse, continuity, missing frames, coordinate scale, conventional speed and displacement metrics, observed and order-null persistence, sequence-dependent persistence, axis contribution, intrinsic axis order, and cue-directed run statistics. These labels match the canonical glossary in main-text Table 1. Design-aware inference is enabled only when the corresponding hierarchical metadata are supplied. The software cannot reconstruct missing biological-replicate identity from pooled trajectory files.

The workflow writes source tables, configuration, seeds, software versions, hashes, numerical checks, and a compressed result archive. Exact analytical nulls avoid permutation loops, but the biological validity of any group contrast still depends on correct experimental labels, units, and hierarchy. Supported statements remain kinematic: the analysis does not identify a unique molecular memory mechanism, prove causal independence from speed, or turn pooled cells into biological replicates.

#### APPENDIX 1 TABLES

Appendix 1—table 1. Complete Step 1–17 computational workflow.

| Step | Purpose | Primary calculation | Frozen result |
|---|---|---|---|
| 1 | Canonical reconstruction | FOV-aware identity, continuity splits, canonical long table | 49,268 raw; 51,450 retained MCF segments |
| 1b | Reconciliation | One-identity-one-longest-segment cohort and accounting | 48,134 primary identities; 117 FOVs |
| 2 | Empirical physics | MSD, local slope, VACF, drift and context tables | Corrected one-hour local slopes >1 |
| 3 | Factorial inference | Repeat/FOV weighting and one-hour contrasts | Positive log-MSD, slope, and VACF interactions |
| 4 | Scale-dependent interaction | Pointwise and simultaneous lag bands | VACF robust 30–140 min |
| 5 | Exact memory decomposition | Direct=activity+memory closure | Raw one-hour interaction memory dominated |
| 6 | Normalized memory dominance | Control-MSD normalization | Direct and memory robust; activity not robust |
| 7 | Fixed-cohort audit | Lag-invariant 24/36/48-point cohorts; LOO FOV | Censoring does not create the effect |
| 8 | Temporal emergence | Matched early-versus-late windows | No reproducible strengthening |
| 9 | Fixed-cohort VACF | 25-point and longer VACF cohorts | VACF robust ≈30–150 min |
| 10 | Temporal-order null | Analytical permutation expectation | Sequence VACF 20–120; covariance 40–100 |
| 11 | Exact memory spectrum | Activity, null memory, sequence memory by lag | Sequence memory robust 120–200 min |
| 12 | Kernel bridge | Pair bands to displacement-memory targets | 20–120→120–200 contribution positive |
| 13 | Kernel enrichment | Opportunity-normalized source/outside densities | Source positive; enrichment unsupported |
| 14 | Evidence freeze | Claims, decision table, source manifest | Primary and unsupported statements frozen |
| 15 | Integrated biophysics | Speed–direction, collective modes, context matching | Directional carrier; negative cross; context robust |
| 16 | Linear GPU model | Local AR flow/residual transitions; 5 scenarios | Full model passed 2/9 targets |
| 17 | Angular GPU HMM | Two-state von Mises model; 6 scenarios | Full model passed 0/8 long-lag targets |

Appendix 1—table 2. Primary raw trajectory identities, retained contiguous segments, and FOV counts by biological repeat.

| Condition | R1 raw / segments | R2 raw / segments | R3 raw / segments | Total raw / segments | FOVs R1,R2,R3 |
|---|---|---|---|---|---|
| shCTRL | 2,915 / 2,985 | 3,573 / 3,691 | 7,731 / 8,264 | 14,219 / 14,940 | 9,9,9 |
| shCTRL + collagen | 1,337 / 1,428 | 2,994 / 3,242 | 5,277 / 5,897 | 9,608 / 10,567 | 10,10,10 |

| Condition | R1 raw / segments | R2 raw / segments | R3 raw / segments | Total raw / segments | FOVs R1,R2,R3 |
|---|---|---|---|---|---|
| shMYO10 | 2,660 / 2,708 | 4,199 / 4,324 | 7,472 / 8,068 | 14,331 / 15,100 | 10,10,10 |
| shMYO10 + collagen | 1,525 / 1,589 | 3,260 / 3,529 | 5,191 / 5,725 | 9,976 / 10,843 | 10,10,10 |

Appendix 1—table 3. Frozen robust intervals and estimates. A dash indicates that the interval, rather than a single scalar, was the primary result.

| Observable | Primary/fixed robust interval | One-hour or target estimate | Key robustness |
|---|---|---|---|
| VACF interaction | 30–150 min | 0.098046 | All 117 FOV omissions positive in all repeats |
| Scaled covariance | 80–150 min | — | Fixed-cohort |
| Sequence-excess VACF | 20–120 min | 0.068217 (0.020851–0.115584) | Order-null has no robust interval |
| Sequence-excess covariance | 40–100 min | 0.072707 (0.021198–0.124216) | All FOV omissions positive |
| Sequence-excess memory | 120–200 min | Target mean 0.181206 | Full fixed cohort |
| 20–120-min source bridge | Target 120–200 min | 0.158036 (0.074057–0.242314) | All FOV omissions positive in all repeats |
| Source density | Target 120–200 min | 0.00086269 (0.00039204–0.00133912) | Repeat signs +,+,+ |
| Source-minus-outside density | No robust interval | 0.000263; interval −0.000819–0.001128 | Repeat signs +,+,− |
| Directional interaction | 50–150 min | — | Only robust speed–direction component |
| Collective flow | 40–60 and 80–140 min; isolated 160, 180 | — | Positive |
| Cell-relative | 70–150 min | — | Positive |
| Flow–cell cross | 10–160 min; isolated 180, 200 | — | Negative |

Appendix 1—table 4. Leave-one-cell-out collective decomposition at 60 min. Values are equal-repeat estimates and simultaneous 95% intervals.

| Component | Mean | Simultaneous 95% interval | R1, R2, R3 | Excludes zero |
|---|---|---|---|---|
| Collective total | 0.115765 | −0.004958 to 0.236487 | 0.081477, 0.036863, 0.228954 | No |
| Shared FOV flow | 0.019295 | 0.000272 to 0.038318 | 0.007931, 0.013362, 0.036592 | Yes |
| Cell-relative | 0.102297 | −0.003085 to 0.207679 | 0.078592, 0.030207, 0.198092 | No |
| Flow–cell cross | −0.005827 | −0.009992 to −0.001663 | −0.005046, −0.006706, −0.005731 | Yes |

Appendix 1—table 5. One-hour VACF interaction under physical-context matching and exclusion sensitivities.

| Scenario | Mean J | R1 | R2 | R3 | All repeats positive |
|---|---|---|---|---|---|
| Unmatched primary | 0.098046 | 0.051766 | 0.062468 | 0.179904 | Yes |
| Occupancy only | 0.100337 | 0.054202 | 0.064457 | 0.182351 | Yes |
| Density only | 0.094738 | 0.049885 | 0.059926 | 0.174404 | Yes |
| Edge only | 0.101629 | 0.056136 | 0.072617 | 0.176133 | Yes |
| Duration only | 0.086973 | 0.040220 | 0.044839 | 0.175859 | Yes |
| Joint nonspatial | 0.096924 | 0.054202 | 0.062968 | 0.173601 | Yes |
| Joint context | 0.103179 | 0.054202 | 0.077090 | 0.178247 | Yes |
| Edge-excluded pairs | 0.098196 | 0.053110 | 0.060413 | 0.181063 | Yes |
| Minimum 48 points | 0.097317 | 0.051948 | 0.059421 | 0.180583 | Yes |

Appendix 1—table 6. Full linear-model frozen predictive validation. Coverage is the fraction of lag points whose observed value lay in the 95% predictive interval.

| Step 16 target | Window (min) | Lag coverage | P(correct sign) | Status |
|---|---|---|---|---|
| VACF | 30–150 | 0.308 | 1.000 | FAIL |
| Sequence VACF | 20–120 | 0.182 | 1.000 | FAIL |
| Sequence covariance | 40–100 | 0.143 | 1.000 | FAIL |
| Directional | 50–150 | 0.273 | 1.000 | FAIL |
| Collective flow | 80–140 | 1.000 | 1.000 | PASS |
| Cell-relative | 70–150 | 0.000 | 1.000 | FAIL |
| Flow–cell cross | 10–160 | 0.000 | 0.4375 | FAIL |
| Sequence memory | 120–200 | 0.889 | 1.000 | PASS |
| Bridge | 120–200 | 0.000 | 1.000 | FAIL |

Appendix 1—table 7. Frequency of the complete expected repeat-sign hierarchy. These frequencies are secondary to quantitative coverage.

| Workflow | Scenario | Complete hierarchy | Frequency |
|---|---|---|---|
| Step 16 linear | Full | 6/128 | 0.0469 |
| Step 16 linear | No interaction | 3/128 | 0.0234 |
| Step 16 linear | One mode | 4/128 | 0.0313 |
| Step 16 linear | No coupling | 12/128 | 0.0938 |
| Step 16 linear | No flow | 2/128 | 0.0156 |
| Step 17 angular | Full | 13/128 | 0.1016 |
| Step 17 angular | No interaction | 6/128 | 0.0469 |
| Step 17 angular | One state | 6/128 | 0.0469 |
| Step 17 angular | No second harmonic | 9/128 | 0.0703 |
| Step 17 angular | No FOV random effects | 13/128 | 0.1016 |
| Step 17 angular | Constant switching | 13/128 | 0.1016 |

Appendix 1—table 8. Held-out local angular likelihood. Two FOVs per repeat-condition group were withheld; random effects were zero for held-out evaluation.

| Angular model | Held-out NLL per emission |
|---|---|
| No FOV random effects | 1.512510 |
| Full | 1.512524 |
| No second harmonic | 1.512761 |
| Constant switching | 1.513265 |
| One state | 1.555430 |

Appendix 1—table 9. Full angular-HMM frozen long-lag validation. The model generally recovered signs but not magnitudes or curve shapes.

| Step 17 target | P(correct sign) | Lag coverage | Status |
|---|---|---|---|
| VACF | 1.000 | 0.000 | FAIL |
| Sequence VACF | 1.000 | 0.000 | FAIL |
| Sequence covariance | 0.992 | 0.000 | FAIL |
| Directional | 1.000 | 0.000 | FAIL |
| Cell-relative | 1.000 | 0.000 | FAIL |
| Flow–cell cross | 0.992 | 0.000 | FAIL |
| Sequence memory | 1.000 | 0.000 | FAIL |
| Bridge | 1.000 | 0.000 | FAIL |

Appendix 1—table 10. Independent external-validation endpoints. Sequence-excess intervals are simultaneous 95% bands at 60 min.

| System | XMLs | Fixed tracks | AUC 20–120 | Sequence excess at 60 [sim. 95%] | Horizon | Serial memory / activity |
|---|---|---|---|---|---|---|
| MDA-MB-231 | 17 | 26,417 | 0.788 | 0.0066 [0.0023, 0.0109] | 70 | −0.741 |
| HUVEC | 18 | 18,641 | 30.917 | 0.3240 [0.3108, 0.3373] | 120 | 2.387 |
| MDCK bulk | 15 | 147,710 | 3.634 | 0.0650 [0.0595, 0.0705] | 60 | 2.037 |
| MDCK edge | 15 | 103,844 | 4.015 | 0.0564 [0.0508, 0.0620] | 60 | 1.755 |

Appendix 1—table 11. Locked haptotaxis contrasts.

| Endpoint | Estimate | 95% interval | Boundary |
|---|---|---|---|
| Lateral order, 20–60 min, 250−mean(20–80) | +0.1643 | 0.1025 to 0.2271 | Short-lag lateral redistribution |
| Gradient-axis contribution, 20-60 min | −0.1182 | −0.2222 to −0.0021 | Opposite-axis shift |
| Total order, 20–60 min | +0.0461 | −0.0542 to 0.1502 | No resolved global increase |
| Intrinsic lateral order, full track | +0.2176 | 0.1274 to 0.3171 | Not only increased lateral activity |
| Fully adjusted intrinsic lateral order | +0.2159 | 0.1277 to 0.3092 | Adjusted for activity, speed, persistence, duration |
| Early minus late lateral contrast | +0.1538 | 0.0745 to 0.2386 | Stronger at short lags |
| 6 h minus 3 h horizon | Unresolved | Includes zero | No discrete onset claim |

Appendix 1—table 12. Primary and balanced PFKL-N702T contrasts.

| Outcome/analysis | Effect | 95% interval |
|---|---|---|
| Mean speed, unadjusted | −0.6769 μm $h^{-1}$ | −1.4239 to 0.1212 |
| Cue FMI, unadjusted | −0.2783 | −0.3575 to −0.2000 |
| Mean up-gradient run, unadjusted | −0.6663 steps | −1.1275 to −0.2039 |
| Cue FMI, HC3 adjusted | −0.2797 | −0.3580 to −0.2014 |
| Mean run, HC3 adjusted | −0.5289 steps | −0.9774 to −0.0803 |
| Cue FMI, 127 matched pairs | −0.2893 | −0.3710 to −0.2053 |
| Mean run, 127 matched pairs | −0.6527 steps | −1.1581 to −0.1641 |
| Cue FMI, overlap weighted | −0.2797 | −0.3573 to −0.2021 |
| Mean run, overlap weighted | −0.5285 steps | −0.9910 to −0.0660 |

Appendix 1—table 13. Generic-order non-inferiority and interpretive boundary.

| Metric | Hedges g | One-sided 95% lower bound | Margin | Conclusion |
|---|---|---|---|---|
| Intrinsic EGF-axis order | +0.1681 | −0.0188 | −0.10 SD | Non-inferior at track level |
| Total serial order | +0.1631 | −0.0233 | −0.10 SD | Non-inferior at track level |
| Cue-sign persistence | +0.1570 | −0.0314 | −0.10 SD | Non-inferior at track level |

### APPENDIX 1 FIGURES

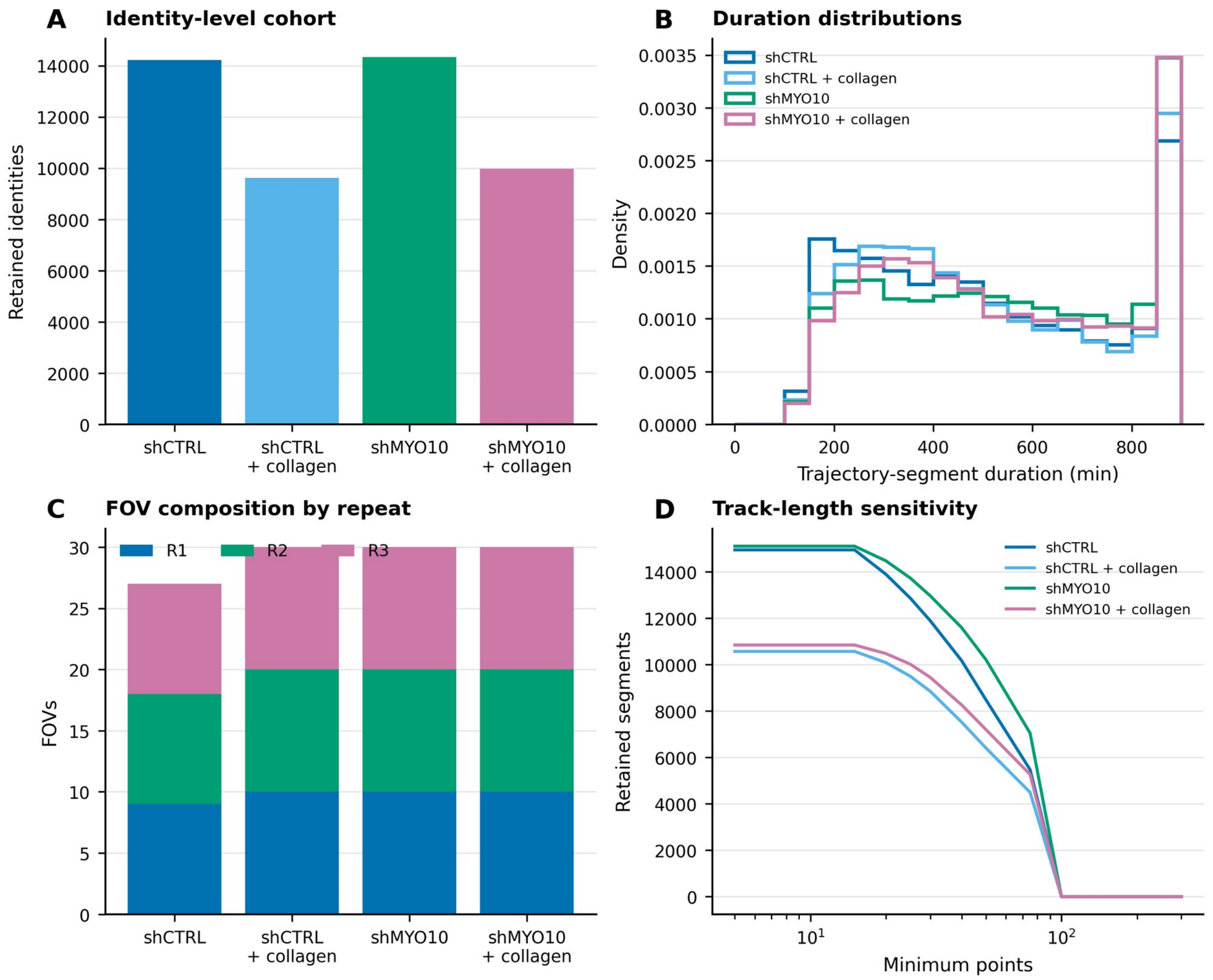


**Appendix 1—figure 1. Corrected cohort audit and provenance. (A) Retained identity-level cohort by condition. (B) Distributions of retained trajectory-segment duration. (C) FOV composition by source-labelled repeat. (D) Number of retained segments as the minimum-point threshold is increased. All counts derive from the corrected FOV-aware reconstruction.**

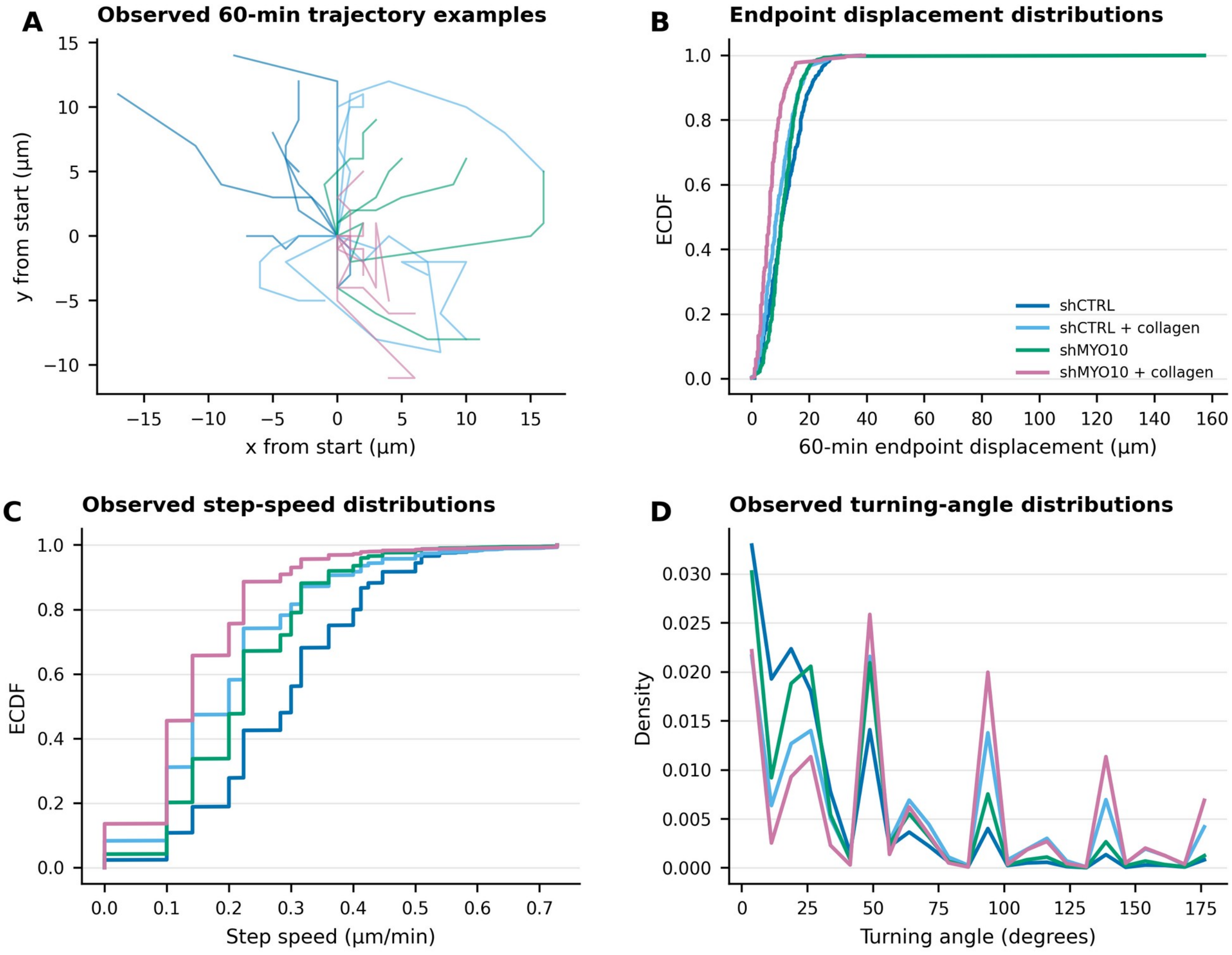


**Appendix 1—figure 2. Genuine trajectory-level biology from the four selected real FOVs. (A) Observed 60-min trajectory examples selected deterministically between the 10th and 90th percentiles of endpoint displacement. (B) Empirical cumulative distributions of 60-min endpoint displacement. (C) Empirical cumulative distributions of observed step speed. (D) Turning-angle distributions. No microscopy background, cell shape, trajectory, or value was generated; all panels use the corrected uploaded coordinates and time points.**

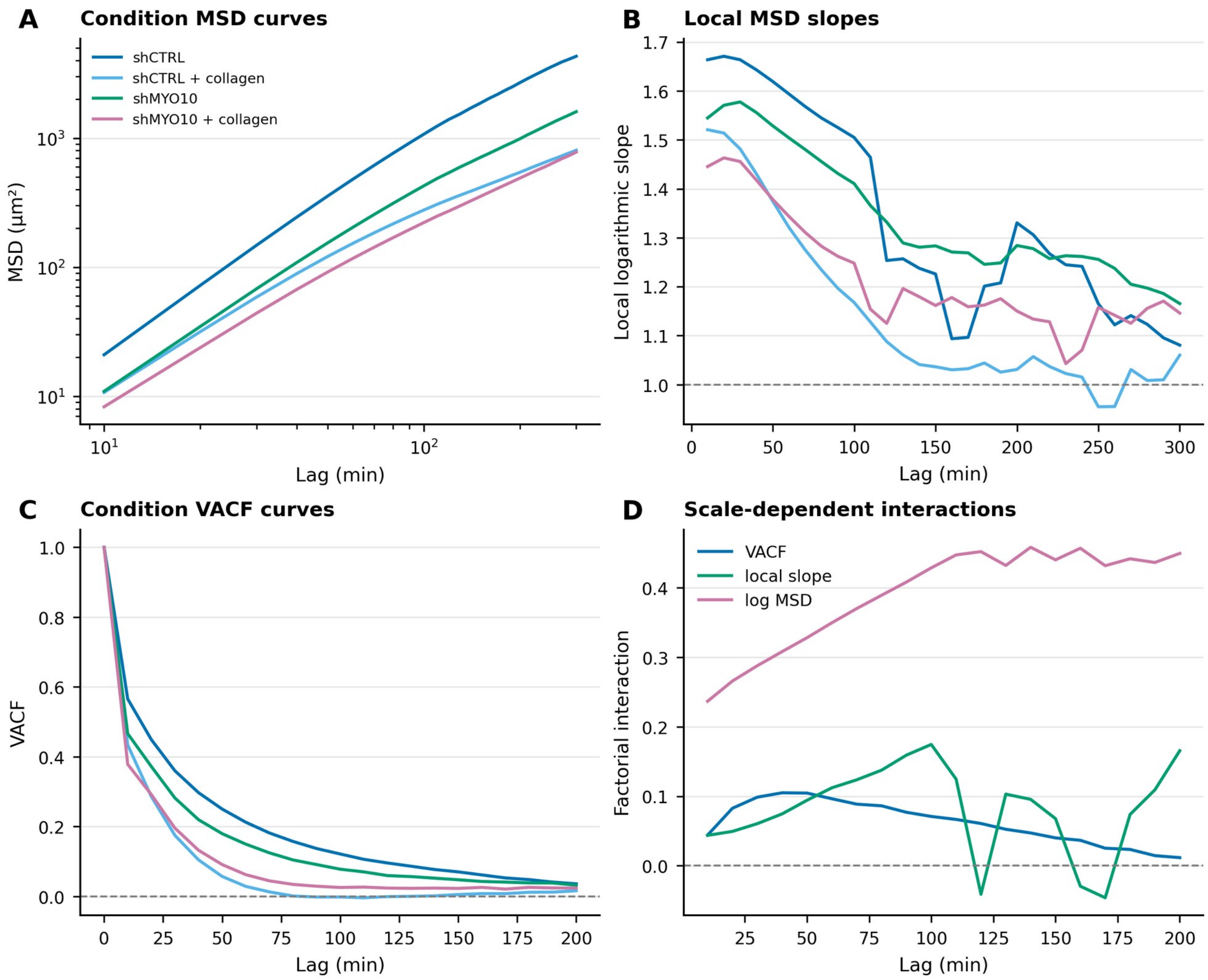


**Appendix 1—figure 3. Scale-dependent corrected empirical physics. (A) Condition-level MSD curves. (B) Local logarithmic MSD slopes. (C) VACF curves. (D) Factorial interactions across lag for VACF, local slope, and log MSD. Curves are aggregated with the frozen FOV/repeat hierarchy; a local slope is not a globally fitted anomalous-diffusion exponent.**

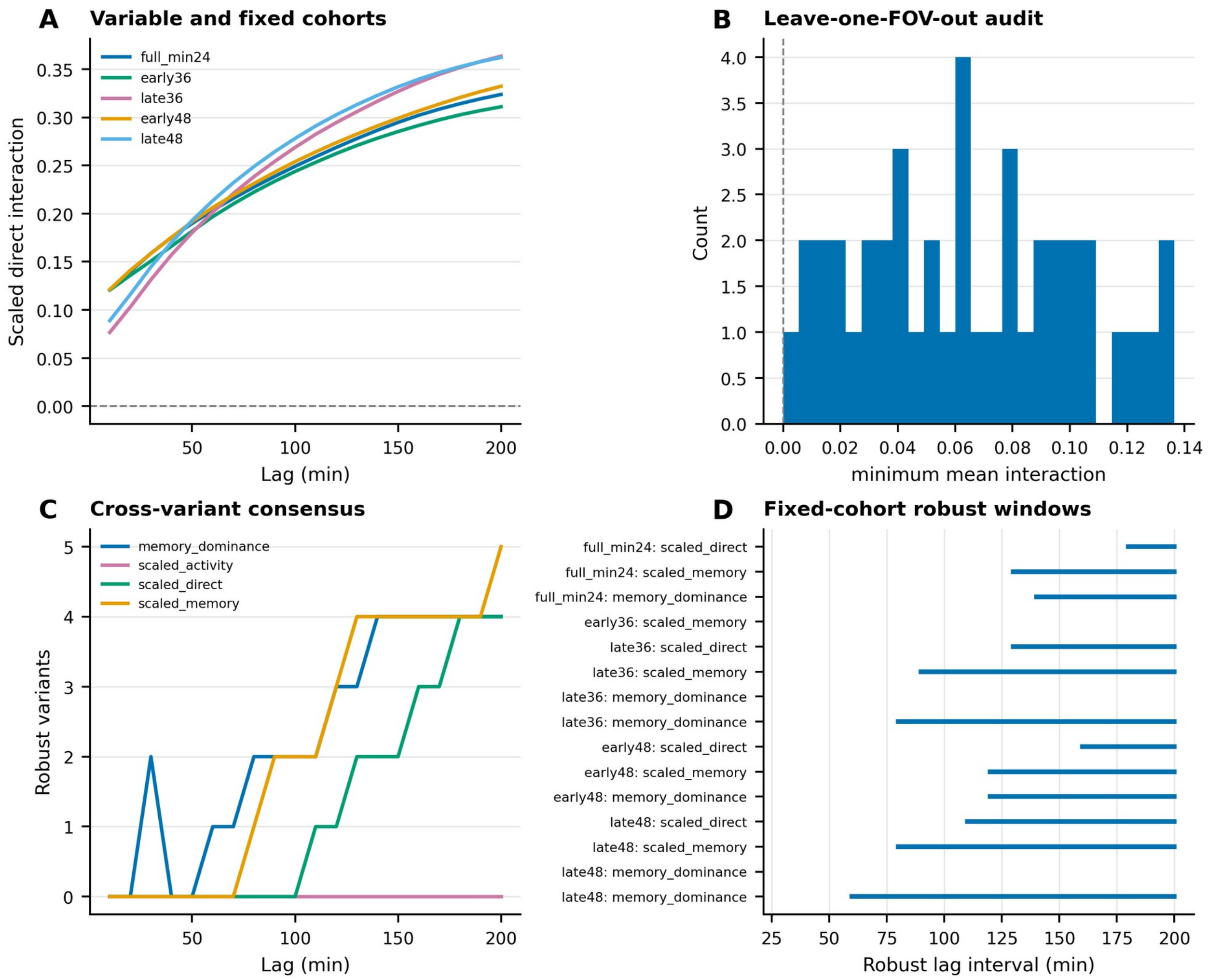


**Appendix 1—figure 4. Fixed-cohort and influence sensitivity. (A) Variable- and fixed-cohort normalized interaction curves. (B) Distribution of the minimum one-hour interaction across leave-one-FOV-out analyses. (C) Cross-variant consensus in the number of robust analyses across lag. (D) Robust lag intervals for prespecified fixed-cohort variants. All 117 FOV omissions retain a positive mean and positive repeat signs at one hour.**

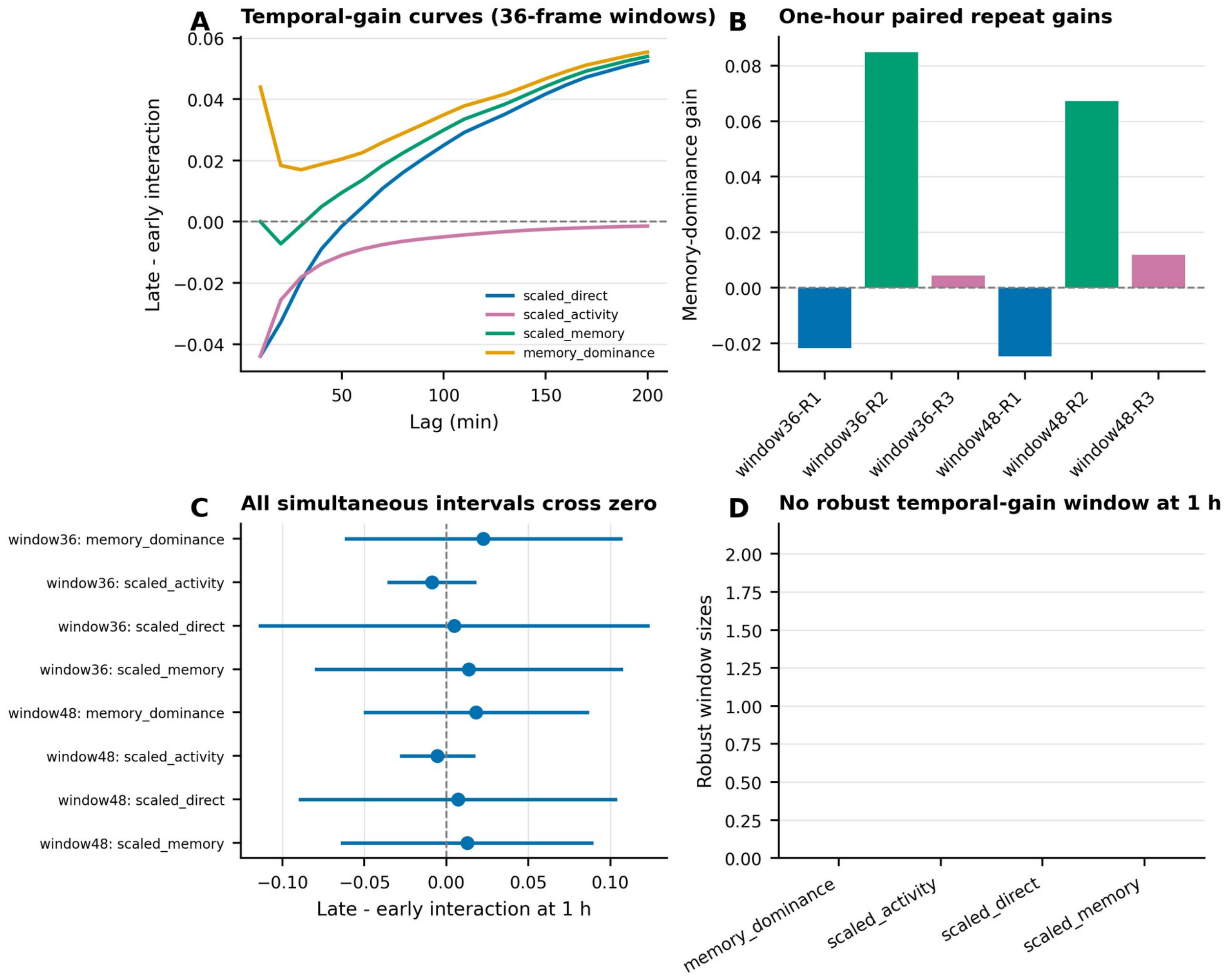


**Appendix 1—figure 5. No reproducible early-to-late strengthening. (A) Late-minus-early interaction curves for matched 36-frame windows. (B) Repeat-level one-hour paired gains for 36- and 48-frame windows. (C) Simultaneous intervals for all paired gain metrics cross zero. (D) Number of robust temporal-gain windows at one hour; no prespecified metric has a robust window.**

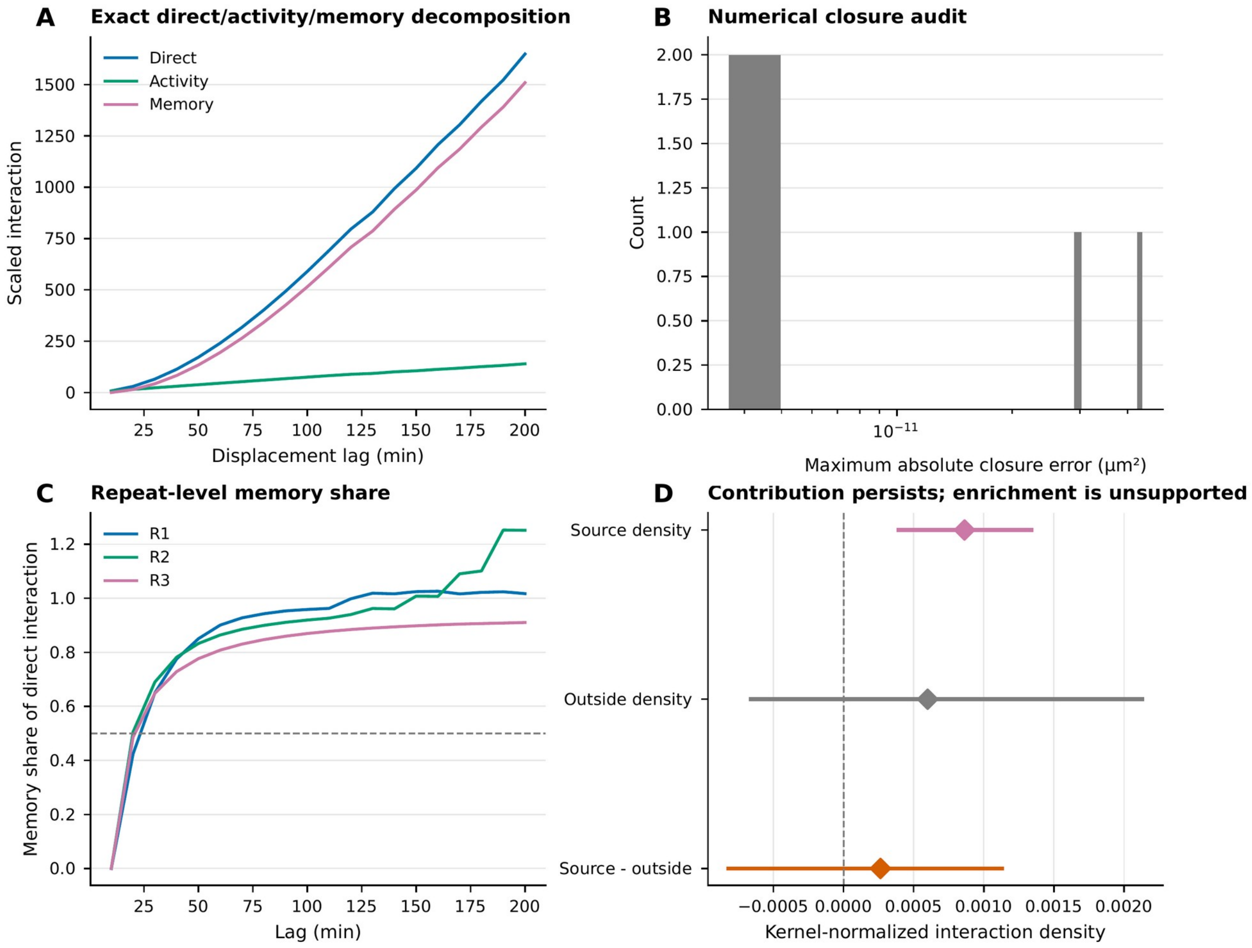


**Appendix 1—figure 6. Exact closure and kernel-opportunity normalization. (A) Direct interaction and its exact activity-plus-memory reconstruction across lag. (B) Numerical closure errors, demonstrating machine-precision implementation of the algebraic identities. (C) Repeat-level memory share across lag. (D) Kernel-normalized source density, outside density, and source-minus-outside contrast. The source density remains positive, whereas enrichment relative to outside-band opportunity is not supported.**

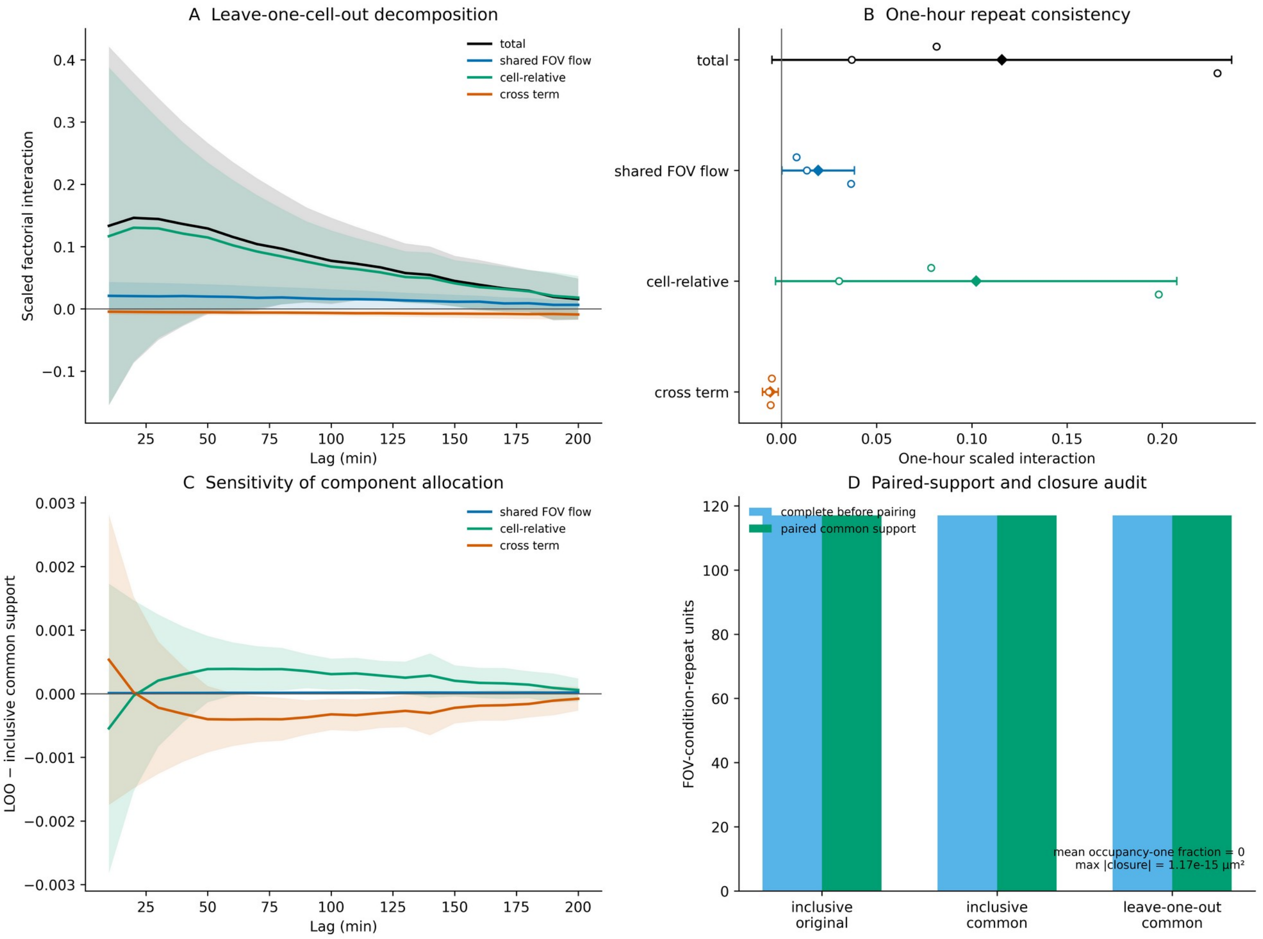


**Appendix 1—figure 7. Leave-one-cell-out collective-flow sensitivity. (A) Lag-dependent leave-one-cell-out decomposition into total, shared FOV flow, cell-relative, and cross components with simultaneous 95% bands. (B) One-hour equal-repeat means, simultaneous intervals, and R1-R3 estimates. (C) Leave-one-cell-out minus inclusive-common allocation differences on identical support. The total is exactly invariant and component differences are small. (D) Paired-support and closure audit. All 117 FOV-condition-repeat units retain support, the occupancy-one fraction is zero, and closure is at machine precision.**

**Appendix 1—figure 8. Fractional active-memory model diagnostics from posterior predictive validation**

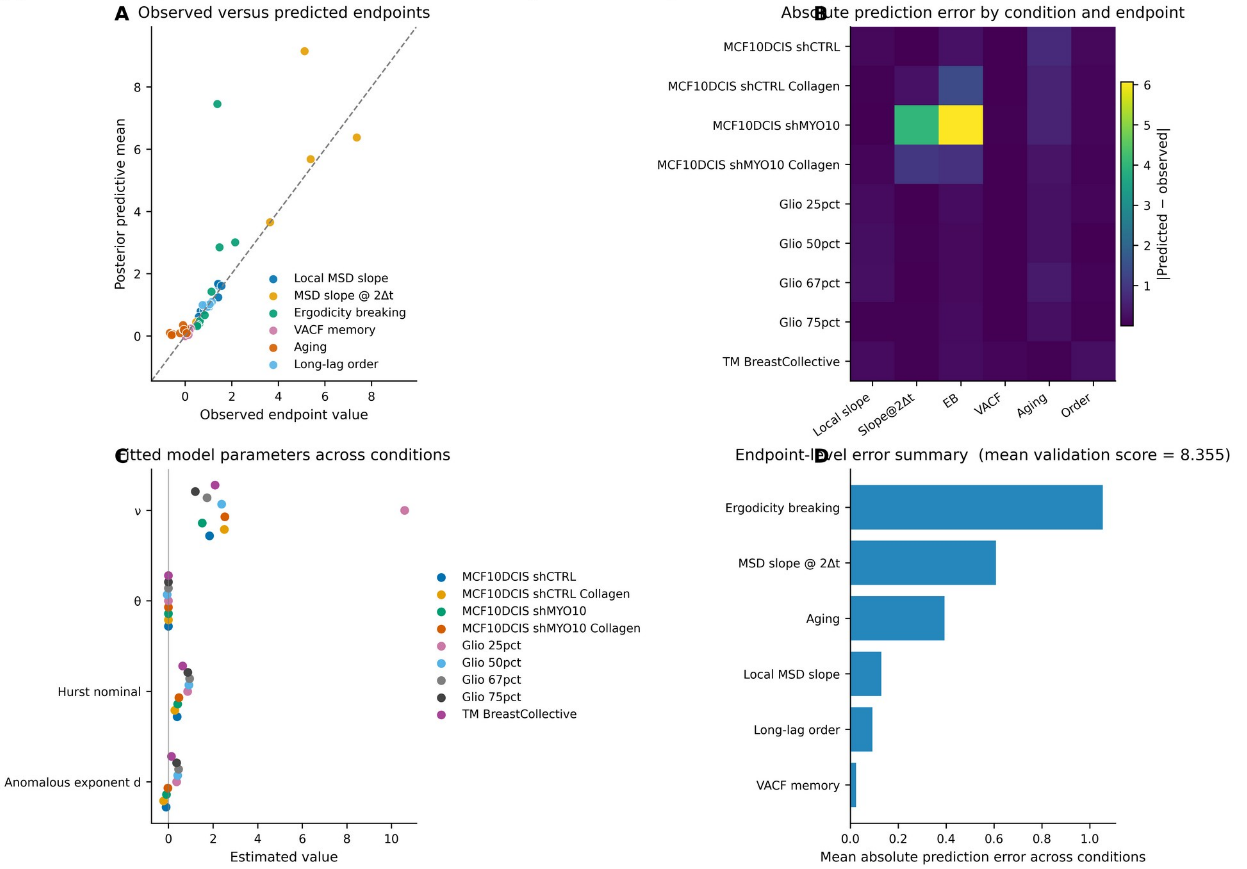


Real-data source: Step8_FractionalModelZoo/posterior_predictive_validation.csv. Panels summarize posterior predictive performance across nine calibration conditions without synthetic redraw of trajectories or endpoints.

**Appendix 1—figure 8. Fractional active-memory model posterior-predictive diagnostics. (A) Observed endpoint values versus posterior-predictive means across nine calibration conditions for local MSD slope, the two-step MSD slope, ergodicity breaking, VACF, aging, and long-lag order; the dashed line is identity. (B) Absolute prediction-error matrix by condition and endpoint. (C) Fitted parameter estimates across conditions. (D) Mean absolute prediction error by endpoint; the title reports the mean validation score. All values are plotted directly from the archived posterior_predictive_validation.csv table.**

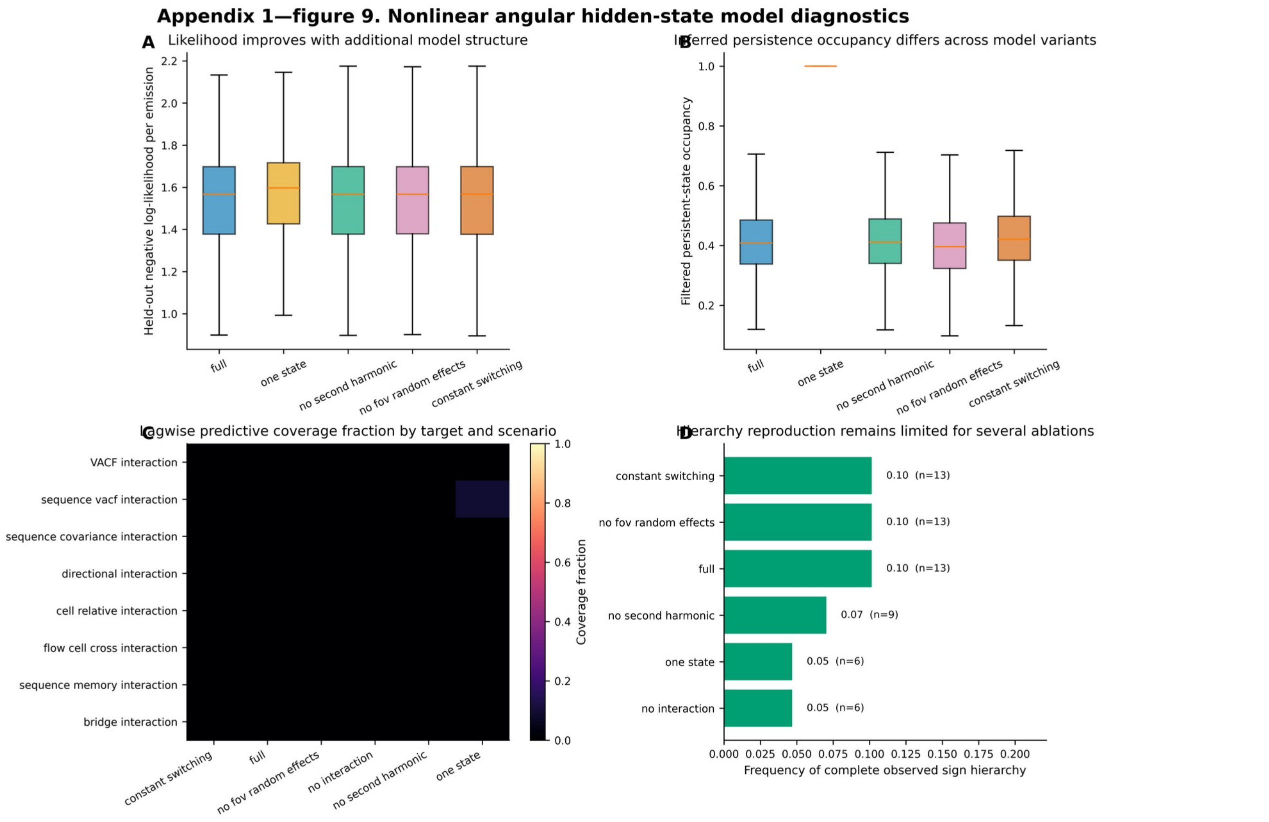


Real-data sources: heldout_angular_likelihood.csv, angular_hmm_validation_matrix.csv, and angular_hmm_hierarchy_frequencies.csv. No synthetic replacement of targets was used; all plotted summaries derive directly from the archived model outputs.

**Appendix 1—figure 9. Nonlinear angular hidden-state model diagnostics. (A) Held-out negative log-likelihood per angular emission across model variants; lower values are better. (B) Filtered persistent-state occupancy across variants. (C) Lagwise predictive coverage fraction by frozen target and simulation scenario. (D) Frequency of reproducing the complete observed sign hierarchy. Two-state local heterogeneity improves held-out likelihood over one state, but the tested models do not reproduce the frozen long-lag hierarchy quantitatively.**

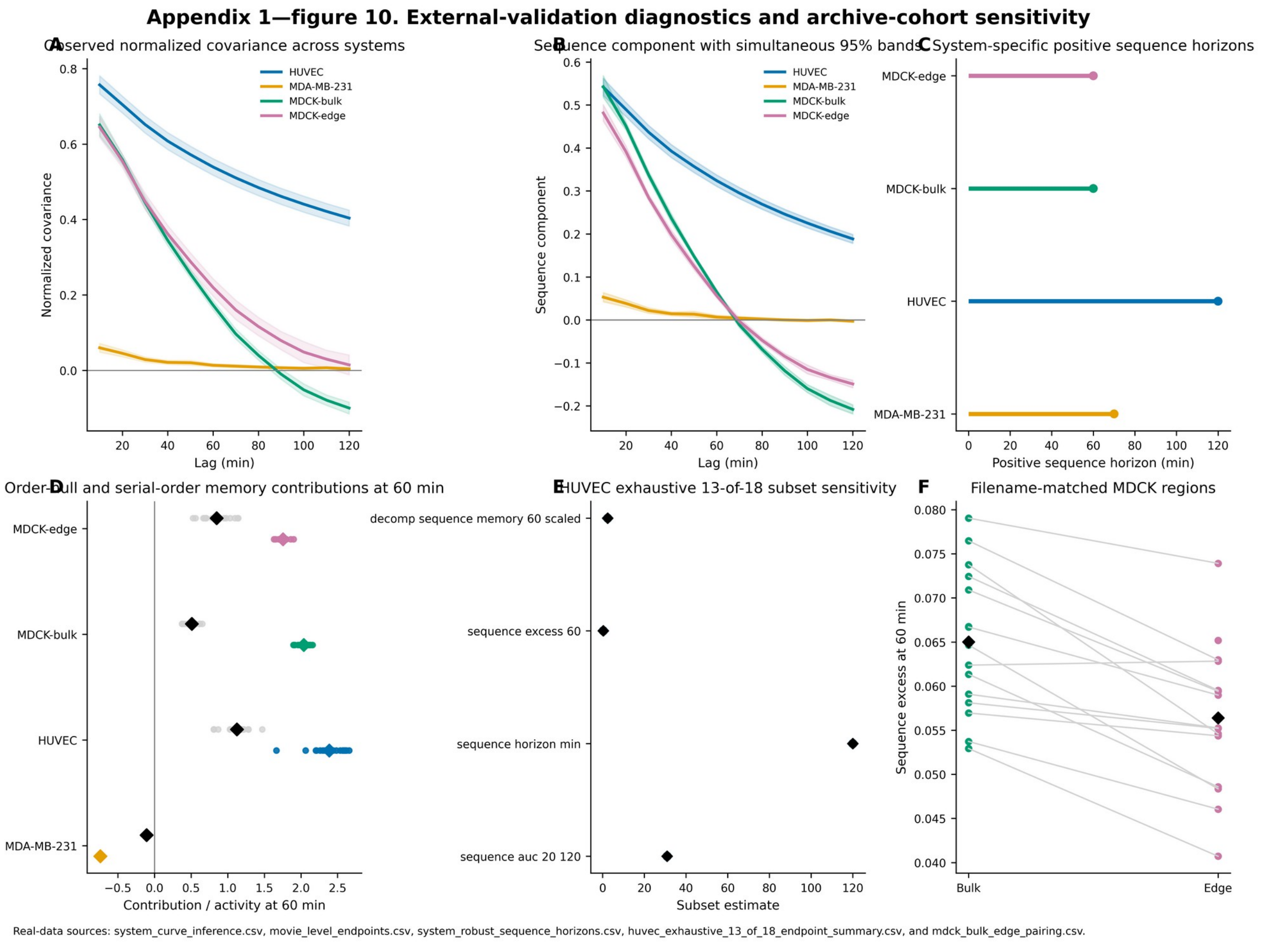


**Appendix 1—figure 10. External-validation diagnostics and archive-cohort sensitivity. (A) Observed normalized covariance across systems. (B) Sequence component across lag with simultaneous 95% bands, including the late-lag sign reversal in MDCK. (C) Simultaneously positive sequence horizon by system. (D) Movie-level order-null and serial-order memory contributions to exact 60-min displacement. (E) Exhaustive HUVEC 13-of-18 subset sensitivity: thin intervals span the full subset range, thick intervals the central 95%, open circles the subset medians, and black diamonds the all-18 estimates. (F) Sequence component at 60 min in filename-matched MDCK bulk and edge regions. Cross-system contrasts remain descriptive because biology and acquisition differ together.**

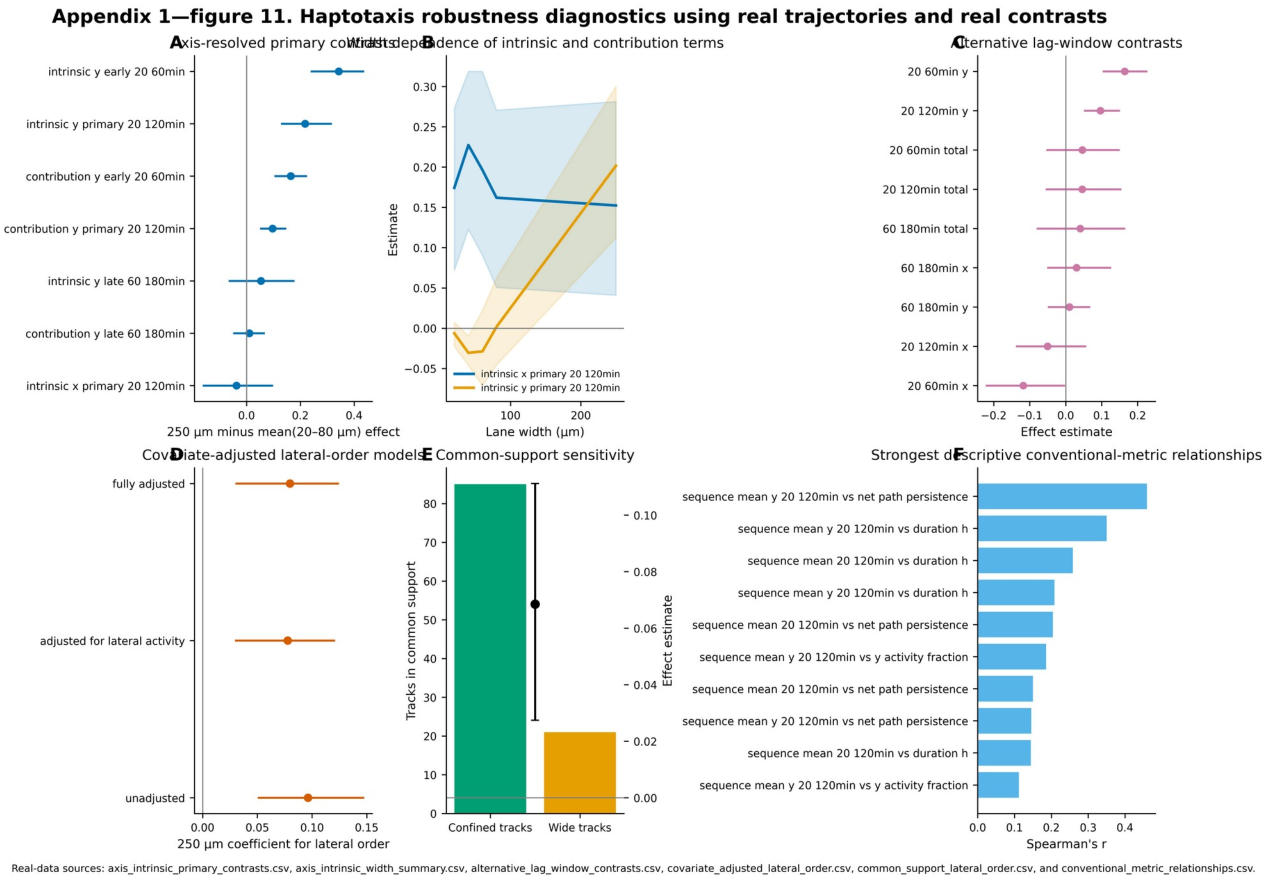


**Appendix 1—figure 11. Haptotaxis robustness diagnostics. (A) Primary wide-minus-confined axis-resolved contrasts with hierarchical 95% intervals. (B) Width dependence of intrinsic-axis order and axis-contribution terms. (C) Alternative lag-window contrasts. (D) Covariate-adjusted lateral-order models. (E) Common-support sensitivity, showing retained confined and wide tracks together with the effect estimate. (F) Strongest descriptive relationships with conventional migration metrics. All estimates derive from the validated haptotaxis closure tables; the figure complements rather than duplicates main-text Figure 4.**

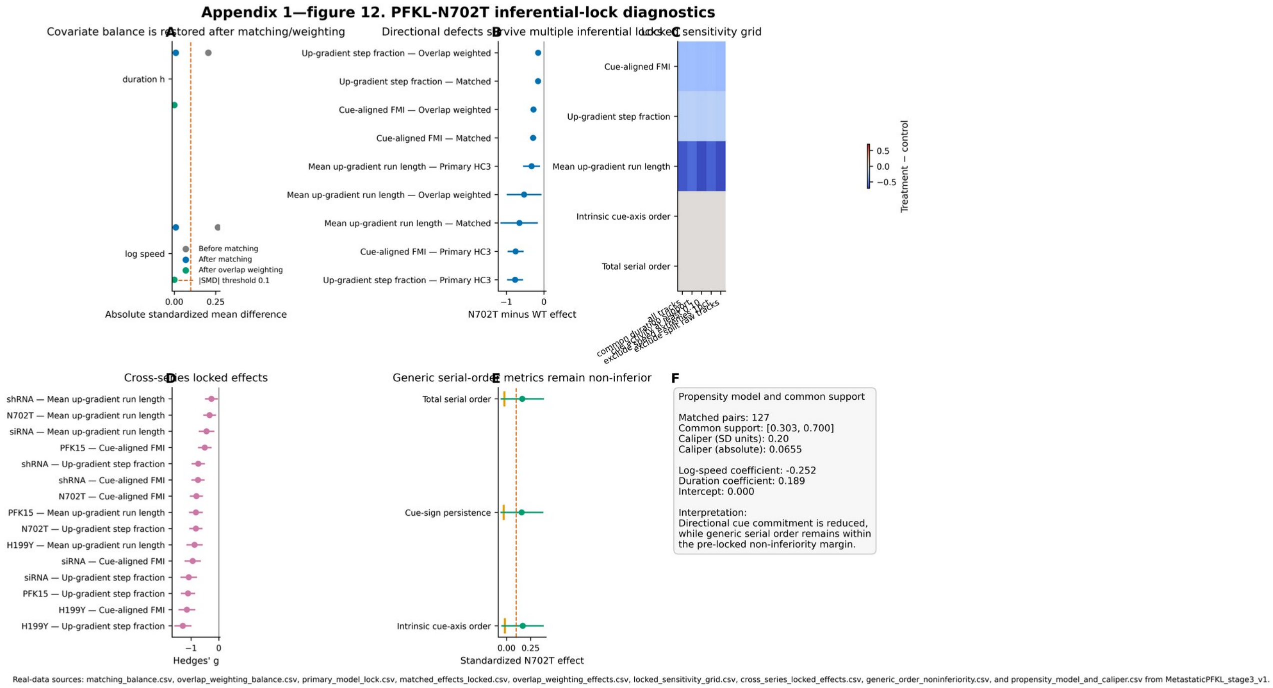


**Appendix 1—figure 12. PFKL-N702T inferential-lock diagnostics. (A) Absolute standardized mean differences before matching, after matching, and after overlap weighting; the dashed line marks |SMD|=0.10. (B) Cue-aligned directional effects under primary HC3, propensity-matched, and overlap-weighted analyses. (C) Locked sensitivity grid across filtering choices. (D) Cross-series standardized directional effects for the within-study PFKL perturbation comparisons. (E) Generic serial-order non-inferiority estimates; orange ticks are one-sided 95% lower bounds and the dashed line is the locked −0.10-SD margin. (F) Propensity-model, common-support, and caliper specification. Biological-replicate identity was unavailable in the deposited trajectory labels, so interval-based PFKL statements remain track-level.**
